# Towards a Better Understanding of Reverse-Complement Equivariance for Deep Learning Models in Regulatory Genomics

**DOI:** 10.1101/2020.11.04.368803

**Authors:** Hannah Zhou, Avanti Shrikumar, Anshul Kundaje

## Abstract

Predictive models mapping double-stranded DNA to signals of regulatory activity should, in principle, produce analogous (or “equivariant”) predictions whether the forward strand or its reverse complement (RC) is supplied as input. Unfortunately, standard neural networks can produce highly divergent predictions across strands, even when the training set is augmented with RC sequences. Two strategies have emerged to enforce equivariance: conjoined/“siamese” architectures, and RC parameter sharing or RCPS. However, the connections between the two remain unclear, comparisons to strong baselines are lacking, and neither has been adapted to base-resolution signal profile prediction. Here we extend conjoined & RCPS models to base-resolution signal prediction, and introduce a strong baseline: a standard model (trained with RC data augmentation) that is made conjoined only after training, which we call “post-hoc” conjoined. Through benchmarks on diverse tasks, we find post-hoc conjoined consistently performs best or second-best, surpassed only occasionally by RCPS, and never underperforms conjoined-during-training. We propose an overfitting-based hypothesis for the latter finding, and study it empirically. Despite its theoretical appeal, RCPS shows mediocre performance on several tasks, even though (as we prove) it can represent any solution learned by conjoined models. Our results suggest users interested in RC equivariance should default to post-hoc conjoined as a reliable baseline before exploring RCPS. Finally, we present a unified description of conjoined & RCPS architectures, revealing a broader class of models that gradually interpolate between RCPS and conjoined while maintaining equivariance.

## 1. Introduction

### Explanatory Video

A 22-minute video explaining the paper is available at https://youtu.be/UY1Rmj036Wg.

Despite sharing a largely identical DNA sequence, different cell types in our body exhibit highly distinct behavior due to the cell-type-specific activity of genes, which is determined by a process called *gene regulation*. A key aspect of gene regulation is the cell-type-specific binding of regulatory proteins known as *transcription factors* (TFs) to regions of the genome known as *regulatory elements*. Neural networks -and in particular Convolutional Neural Networks (CNNs) - have emerged as state-of-the-art models for predicting genome-wide signals of cell-type-specific regulatory activity as a function of the underlying DNA sequence (Zhou and Troyanskaya, 2015; Kelley *et al.*, 2015; Quang and Xie, 2019). Over the course of model training, the weights of a CNN’s filters gradually evolve to identify short (~6-20bp) predictive patterns (known as *motifs*) encoded in regulatory DNA sequences that determine the locations and strength of TF binding. Filters in later layers can build on the motifs learned by filters in previous layers in order to recognize higher-order motif syntax patterns that orchestrate the binding of TF protein complexes.

Unfortunately, the standard CNN architectures used for these prediction tasks are based on models developed in the computer vision literature, and thus do not explicitly account for the reverse-complement symmetry of double-stranded regulatory DNA. Specifically, they do not model the fact that complementary base pairing implies that a motif appearing on the forward strand is semantically analogous to one that appears in the reverse-complement (RC) orientation (even though a TF can bind the DNA strands asymmetrically, the presence of a motif in the RC orientation on one strand implies the presence of the motif in the forward orientation on the complementary strand, and thus has equivalent predictive power). For standard CNNs, learning to recognize both the forward and RC versions of a non-palindromic motif is as challenging as learning to recognize two completely different motifs. As a result, such models frequently produce very different predictions depending on whether a strand is supplied in the forward vs. the RC orientation, even when the training dataset is augmented to contain reverse-complements (Shrikumar *et al.*, 2017). Differing predictions erode confidence in model interpretation, as they could imply that motifs are missed on either the forward or RC strands.

Early work in deep learning for genomics handled this by combining model predictions across forward and RC versions of the input - for example, DeepBind (Alipanahi *et al.*, 2015) took the maximum prediction across both strands, while FactorNet (Quang and Xie, 2019) took the average across strands. Such architectures are sometimes called conjoined a.k.a. “siamese” architectures (Bartoszewicz *et al.*, 2020). While the terms “conjoined”/“siamese” usually imply that the representation merging was performed during both training and testing time (as was done in Alipanahi *et al.* (2015), Quang and Xie (2019) and *Bartoszewicz et al.* (2020)), one can also take a model that was trained without representation merging and perform the merging only during testing time. Although it has not been described as such, merging of representations during test-time is equivalent to converting a standard model to a conjoined one post-training (post-hoc). To date, no work has investigated whether trained conjoined models provide any benefits over post-hoc conjoined models when the training dataset is augmented with reverse-complements. Note that when the form of representation merging is “averaging”, a post-hoc conjoined model is essentially ensembling the model predictions across the forward and RC strands. The conjoined architecture used in this work for binary prediction tasks is illustrated in **Fig. 1**.

**Figure 1.**
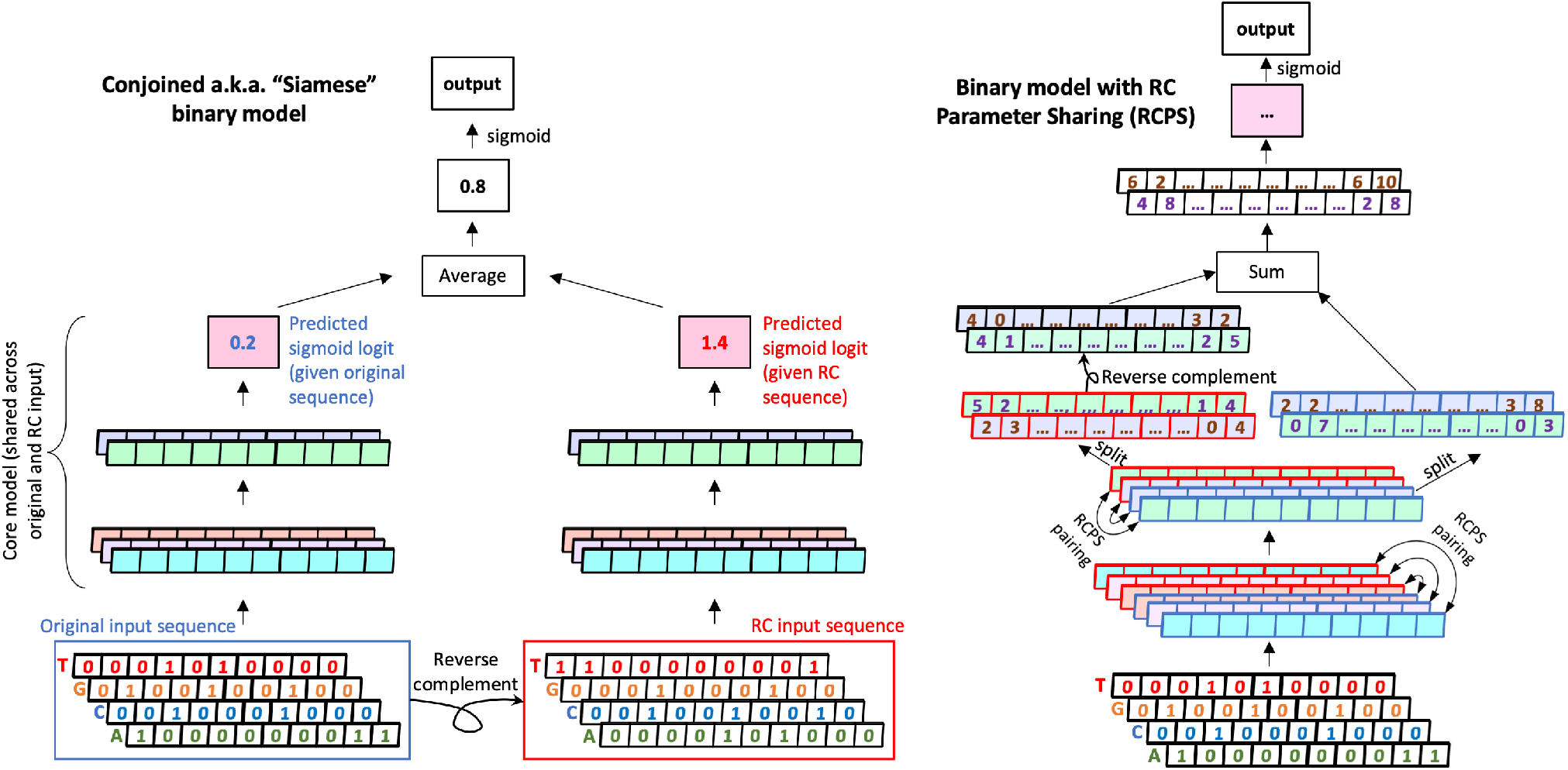
Architecture of conjoined models and RCPS models used in this work for binary prediction. For the conjoined model (**left**), an identical “core model” is applied to both the forward and reverse-complement input sequence, and the predictions of the two branches are averaged prior to applying the final sigmoid nonlinearity. Cells with similar shading represent neurons with shared parameters. In the case of the RCPS model (**right**), the weights of the “forward” filters are paired with corresponding “RC” filters via RCPS (**Fig. 2**), such that each forward filter recognizes the reverse-complement of whatever pattern its matching RC filter recognizes. Once again, cells with similar shading represent neurons with shared parameters - however, blue outlines denote the “forward” versions of a filter, while red outlines denote the corresponding RC versions. The representations learned by the “forward” and “RC” filters are merged after the last convolutional layer by applying the “reverse complement” operation (i.e. flipping along both the length axis and channels axis) to the RC filters and summing. For both the Conjoined and RCPS models, the last convolutional layer is followed by a local maxpooling operation (not depicted); this maxpooling operation is applied before the forward and RCPS filters are summed.

A notable drawback of conjoined architectures is that forward and RC versions of a non-palindromic motif must still be learned as though they are two completely separate motifs - i.e. the model has no explicit knowledge of DNA double-strandedness. While it is true that the conjoined model scans both the forward and RC version of the input sequence, it is possible for a single input sequence to contain multiple motifs where some motifs may be in the forward orientation and others may be in the RC orientation; a model that only recognizes one orientation for each motif will never correctly identify all the motifs present irrespective of whether it is looking at the forward or the RC input sequence. To address this, Shrikumar *et al.* (2017) proposed RC parameter sharing (RCPS). In RCPS, the weights of every convolutional filter are paired with those of a corresponding “RC” version of the filter such that the “RC” filter will recognize the reverse-complement of whatever pattern the forward filter recognizes. This weight sharing is illustrated in **Fig 2**, and the RCPS architecture that we use in this work for binary prediction tasks is illustrated in **Fig. 1**. This idea of RCPS has been employed in several subsequent works: Brown and Lunter (2019) extended RCPS to models with dropout and applied it to predict recombination hotspots, Bartoszewicz *et al.* (2020) applied it to predict the pathogenic potential of novel DNA, and *Onimaru et al.* (2020) used RCPS-like concepts in layers that they refer to as FRSS (Forward and Reverse Sequence Scan). Nevertheless, to date, there has not been a systematic benchmark of RCPS against trained or post-hoc conjoined models that have similar architectures.

**Figure 2.**
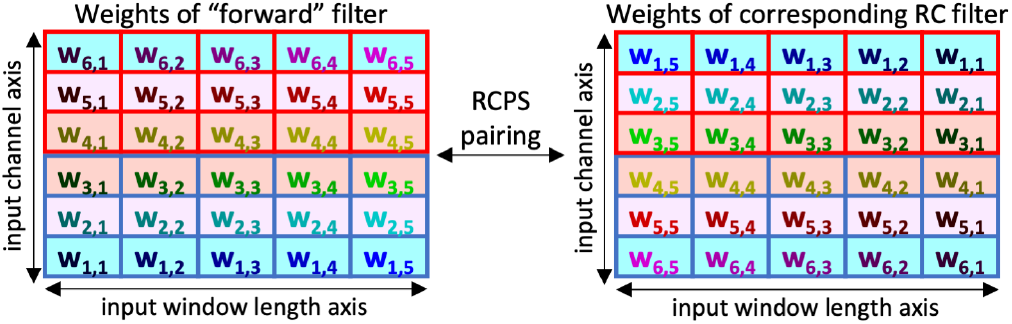
RCPS pairing of convolutional filter weights (Shrikumar *et al.*, 2017). In RCPS pairing, the weights of an RC filter are designed to recognize the reverse-complement of the pattern that the matching forward filter recognizes. RCPS presumes that the “reverse-complement” of the input can be obtained by reversing both the length and channel axes. In the figure, table cells with similar shading represent input channels that are paired; blue out-lines denote the “forward” version of an input channel, while red outlines denote the corresponding RC version. Entries in the table cell denote the filter weights; weights that are paired between the forward and RC filters are given the same text color and subscripts. Note that when the input is a one-hot encoded sequence, the “channel” axis is assumed to encode ACGT in that order; this results in “A & T” and “C & G” being treated as RC-paired input channels.

A third limitation of the existing literature on RC architectures is that they have not been extended to the task of single base-pair resolution signal profile prediction, which has demonstrated immense potential to learn high-resolution, higher-order syntax patterns of TF binding (Avsec *et al.*, 2020). The existing state-of-the-art model for profile prediction is the BPNet architecture (Avsec *et al.*, 2020), which predicts the shape of the observed signal (in the form of a probability distribution over a 1kbp interval) at base-pair resolution using both DNA sequence and a “control” (experimental bias) signal track as input. Separate predictions are made for the forward and reverse strands; this separation enables modeling of the asymmetric “strand shift” found at TF binding sites in profiles obtained from TF binding assays such as ChIP-seq (chromatin immunoprecipitation followed by sequencing) and ChIP-nexus/exo. Extending BPNet architectures to produce analogous (or “equivariant”) predictions for RC sequences would need to handle reverse complements at multiple stages of the input and output (**Fig. 3**).

**Figure 3.**
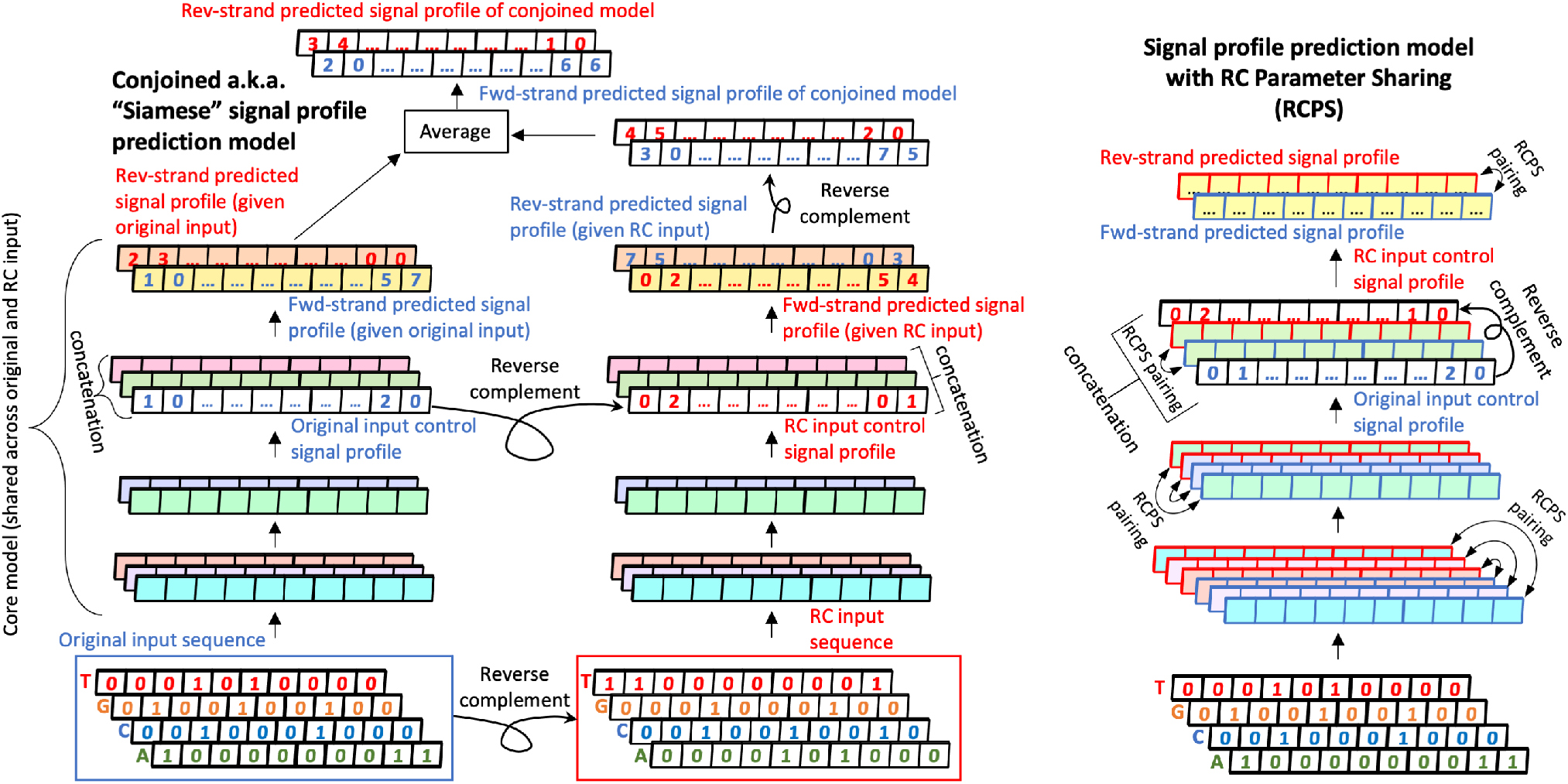
Guaranteeing RC equivariance for the task of base-pair-resolution signal profile prediction. Corresponding architectures for binary output models are in **Fig. 1**. In the standard BPNet profile prediction architecture, which is fully convolutional, the input control signal profile is concatenated as an extra channel to the activations of an intermediate convolutional layer, and separate predictions are made for positive and negative strands. **Left:** conjoined BPNet architecture for profile prediction. Cells with similar shading represent convolutional neurons with shared parameters. Unlike previously-proposed conjoined architectures for binary tasks, the model output on forward and RC inputs cannot simply be averaged; due to the strand-specific profile predictions, the output on the RC input needs to be reverse-complemented to be compatible with predictions on the forward input. **Right:** RCPS architecture for profile prediction. Cells with similar shading represent convolutional neurons with shared parameters; blue outlines denote the “forward” versions of a filter, while red outlines denote the corresponding RC versions. Unlike the RCPS architectures used in binary models, the forward and reverse filters are never collapsed into a single representation; this allows the model to make strand-specific predictions. To maintain RC equivariance, the input “control track” signal profile must be appended to both ends of the convolutional filter stack (once in the forward orientation and once in the RC orientation). Also note that, in the RCPS formulation, the last two convolutional layers only specify the weights for one filter; this is because these layers are intended to contain exactly two channels (one for each DNA strand), and specifying the weights for one filter will result in two output channels (due to RCPS pairing). For RCPS convolutional layers besides below last two layers, weights are specified for the same numbers of filters as in the standard models.

## 2. Our Contributions

1. We devise an extension of Conjoined and RCPS architectures to base-pair-resolution signal profile prediction
2. We establish a strong baseline of post-hoc conjoined models and conduct systematic benchmarks on diverse tasks
3. We find that post-hoc conjoined models consistently perform as well as or better than trained conjoined models, and develop a mathematical intuition for why; we find empirical support for this by studying train vs. test-set performance
4. We prove that the representational capacity of the RCPS models encompasses that of a conjoined model, but nevertheless observe that the RCPS underperforms relative to post-hoc conjoined models on some tasks. We find that this disparity is not easily attributed to overfitting of RCPS, hinting at optimization difficulties.
5. We develop a unified description of conjoined and RCPS architectures, including a novel class of architectures that incrementally interpolates between a fully-conjoined and a full RCPS architecture while maintaining RC equivariance.

## 3. Methods

Full details on the architectures and datasets are provided in supplementary **Sec. C**. In brief: we created two simulated datasets consisting of synthetic DNA sequences of lengths 200bp and 1kbp respectively, that contained motif instances sampled from 3 different TF motif models (Position Weight Matrices i.e. PWMs). Multi-task CNNs (with 3 binary output tasks) were trained to predict whether a given sequence contained instances of a particular motif. We also used genome-wide binarized TF-ChIP-seq data for Max, Ctcf and Spi1 in the GM12878 lymphoblastoid cell-line (Shrikumar *et al.*, 2017). In these data, the positive set contained 1kbp sequences centered on high-confidence TF ChIP-seq peaks, and the negative set contained 1kbp sequences centered on chromatin accessible sites (DNase-seq peaks) in the same cell-lines that do not overlap any TF ChIP-seq peaks. Single-task binary output CNNs were trained on these data. For the base-pair-resolution signal profile prediction, we used genome-wide ChIP-nexus profiles of four TFs - Oct4, Sox2, Nanog and Klf4 in mouse embryonic stem cells (Avsec *et al.*, 2020). BPNet-style models were trained with a multinomial loss to predict the distribution of reads in 1kbp regions for each of the two strands within ChIP-nexus peaks. Separate models were trained for each TF. The conjoined and RCPS architectures used are illustrated in **Fig. 1** and **Fig. 3**. Please refer to https://github.com/hannahgz/BenchmarkRCStrategies for code and links to the data.

### 3.1. A note on the number of filters in RCPS

In the case of the RCPS architectures, each filter has a reverse-complement counterpart that is created at runtime (via weight sharing), which increases the representational capacity of the model. Thus, the “effective” number of filters in the RCPS model could be considered to be twice the number in the standard models. For thoroughness, we also ran benchmarks where the RCPS models were specified to have half the number of filters, such that the “effective” number of filters would be comparable to that in the standard models. However, we found that this consistently decreased the performance of RCPS on all the tasks we evaluated (**Sec. D.5**). Also note that our default setup, where the “effective” number of filters we use for the RCPS model is twice the number in the standard model, is necessary for the proof showing that our RCPS models can represent any solution learned by the corresponding conjoined models; this is because our proof relies on mapping the activations of the conjoined model on the RC input sequence to the activations of the “RC filters” in the RCPS model.

### 3.2. A note on RC data augmentation

Standard architectures (that are not RC-equivariant by design) can be trained with or without data augmentation. When RC data augmentation was enabled, it was implemented by extending each training batch with the reverse complements of the original inputs, which effectively doubles the batch size. A natural question that the reader might have is why we do not train RC-equivariant architectures (i.e. Conjoined and RCPS architectures) with data augmentation. Due to the symmetries present in these architectures, the gradient update performed on a sequence in the forward orientation is *identical* to the gradient update performed on the reverse-complement; thus, if we were to train RC-equivariant architectures with data augmentation, it would be equivalent to duplicating the examples in each batch.

## 4. Results

### 4.1. Post-hoc conjoined models outperform trained conjoined models

Across all datasets, we found that post-hoc conjoined architectures (when trained with RC data augmentation) consistently perform as well as or better than trained conjoined architectures. In fact, post-hoc conjoined models consistently achieved the best performance on the profile prediction tasks (**Fig. 4**), even when their trained conjoined counterparts failed to significantly outerperform standard models trained with data augmentation.

We developed a hypothesis to explain this behavior, which we illustrate first with a theoretical argument: consider a standard (non-conjoined) model *f* that gives a too-high prediction for an input sequence *S* and a too-low prediction on the reverse-complement *S′*. If the model were trained with data augmentation, both *S* and *S′* would be separate examples in the same batch, and the gradient descent update would raise *f* (*S′*) while lowering *f* (*S*). By contrast, when training with the conjoined version of *f*, there would be no separate loss computed for *f* (*S*) and *f* (*S′*); only (*f* (*S′*) + *f* (*S*))*/*2 would be compared to the true value. Thus, if (*f* (*S′*) + *f* (*S*))*/*2 were close to the true value, the gradient descent update for the conjoined model would not change *f* (*S′*) or *f* (*S*), even though the model has not learned the true value in either case. Models that are conjoined during training may therefore overfit to the training set and fail to converge to the most generalizable solution - for example, the may latch on to artifactual signal on *S* while failing to recognize true motifs on *S′*.

**Figure 4.**
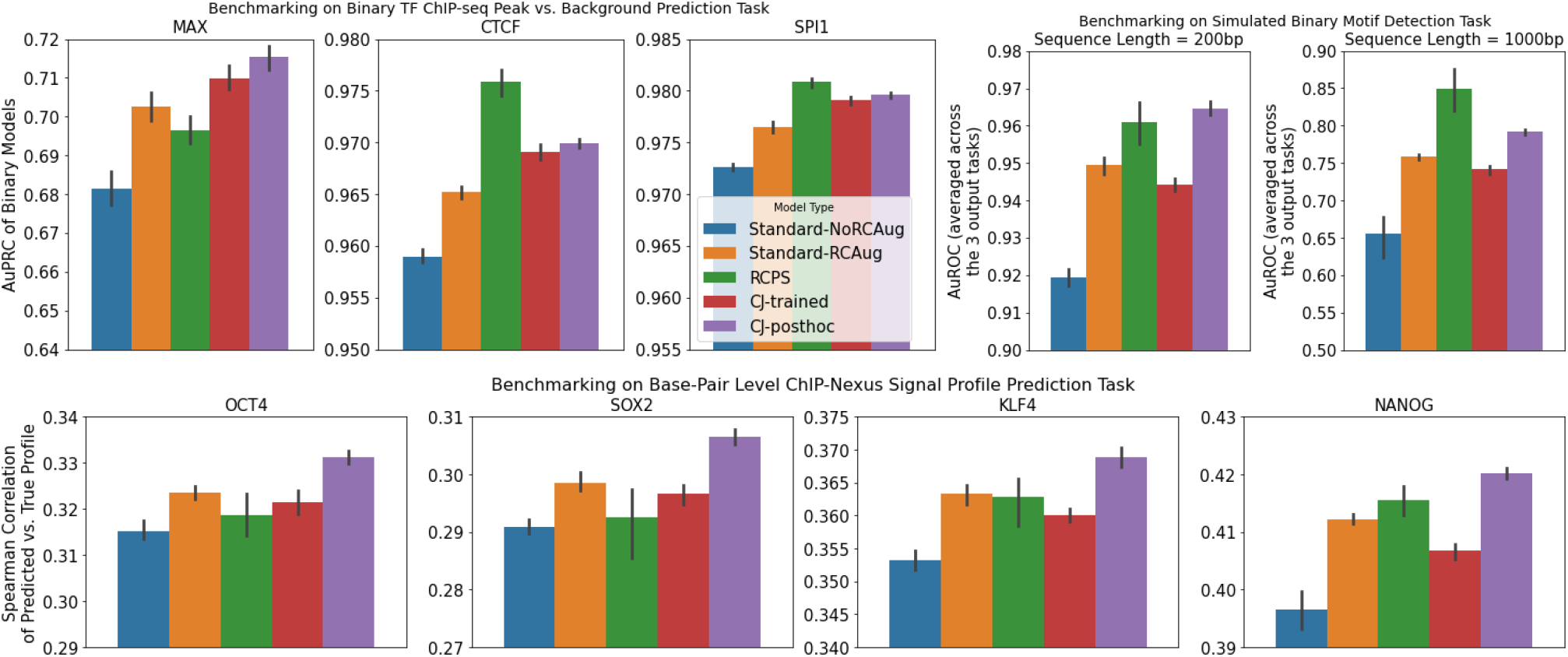
Benchmarking models on binary TF ChIP-seq peak prediction, simulated binary classification, and base-pair-resolution signal profile prediction tasks. Bar heights represent the average performance over 10 random seeds, and error bars represent the 95% confidence intervals for the mean generated by seaborn (Waskom *et al.*, 2017) using 1000 bootstrapped samples. “Standard-RCAug” and “Standard-noRCAug” are standard models trained with and without RC data augmentation. RCPS denotes RC Parameter Sharing. CJ-trained are models that were conjoined during both training and test time. CJ-posthoc are standard models trained with RC data augmentation that were converted to conjoined models only after training. CJ-posthoc consistently performs as well or better than all other methods on all tasks except CTCF ChIP-seq, SPI1 ChIP-seq, and 1000bp simulated (all of which involve binary prediction). On these tasks, RCPS does best - however, RCPS does not perform significantly better than StandardRCAug on all profile prediction tasks and the Max binary task (as per overlapping confidence intervals). For the binary TF ChIP-seq models, the training hyperparameters were set to the tested combination that gave the best overall AuPRCs: 8000 (rather than 4000) maximum training iterations, AuROC (rather than model loss) as the metric used to choose the best validation-set epoch, and upsampling (rather than upweighting) of positives examples to achieve a 1:4 class ratio during training (see **Sec. D.3** for more details, including a reporting of the results with AuROC rather than AuPRC as the performance metric). For the simulated datasets, we report AuROC (rather than AuPRC) as the main performance metric as the dataset was not imbalanced. As with the binary TF ChIP-seq models, and consistent with Shrikumar *et al.* (2017), the best validation-set epoch for the binary simulated models was selected using AuROC (rather than model loss). For the profile prediction models, we report the Spearman correlation of the true signal profile vs. the predicted signal profile as the metric; plots showing different performance metrics for profile prediction are in **Fig. D.6**, and the profile metrics are explained in **Sec. C.2.3**.

However, alternative explanations for poor performance are also possible; for example, it is known that multitasked deep learning models can be challenging to optimize because the gradients from different tasks may conflict with each other during training (Yu *et al.*, 2020); perhaps the two branches of the conjoined model may similarly result in conflicting gradients during training. Model initialization can also play a significant role in the performance of a deep learning model, and it is possible that initializations may impact different model architectures differently (note, however, that we train our binary networks with batch normalization, which is thought to reduce sensitivity to initialization (Schilling, 2016)). To investigate our overfitting hypothesis, we compared the training vs. test-set performance of post-hoc conjoined models to the training vs. test-set performance of the trained conjoined models (**Fig. D.1**). Consistent with our hypothesis, we found that for the binary TF ChIP-seq and base-pair-resolution signal prediction tasks, the mean training-set performance of the trained conjoined models was comparable to or better than the mean training set performance of the post-hoc conjoined models, even though the mean test-set performance of trained conjoined models was comparable to or worse than the mean test-set performance of post-hoc conjoined models. We can therefore conclude that, for these tasks, a failure to optimize on the training set does not explain the poor performance of trained conjoined models. However, for simulated datasets, we found that the improvement in test-set performance for post-hoc conjoined models was also accompanied by an improvement in training-set performance. We note, however, that the models were all trained with early stopping, and it is therefore possible that the trained conjoined models might have surpassed the post-hoc conjoined models in training set performance if all models were forced to train for the same number of iterations.

### 4.2. RCPS shows inconsistent performance across tasks

While we were able to replicate the performance of RCPS reported by Shrikumar *et al.* (2017) on their binary classification datasets, we made a few interesting observations. First, RCPS on the 200bp simulated sequence dataset did not perform significantly better than data-augmented post-hoc conjoined models (which were not included as a baseline in Shrikumar *et al.* (2017)). Second, as discussed in **Sec. D.3**, RCPS did not perform as well relative to conjoined models and data-augmented standard models on the Max ChIP-seq task when the maximum number of training iterations was increased beyond what Shrikumar *et al.* (2017) used. Third, on base-pair-resolution profile prediction datasets, RCPS *consistently* underperformed relative to post-hoc conjoined models - in fact, the 95% confidence intervals for both RCPS and trained conjoined models either overlapped with or were lower than the confidence interval for data-augmented standard models on these datasets (**Fig. 4**).

These results can be considered particularly surprising given that the solution learned by the post-hoc conjoined models could have equivalently been represented using the RCPS models. The proof of this equivalence is given in **Sec. A**. We thus conclude that the mediocre performance of RCPS is not due to a representational limitation. One hypothesis is that the RCPS models, like the trained conjoined models, may overfit to the training set (**Sec. 4.1**). In fact, the risk of overfitting is arguably greater for RCPS models due to their increased representational capacity (**Sec. 3.1**). As before, we investigated this hypothesis by plotting the training vs. test-set performance of RCPS against that of the conjoined models. However, we found that in the cases where RCPS underperforms relative to post-hoc conjoined models on the held-out set, the training-set performance of the RCPS models was never significantly better than that of the post-hoc conjoined models (**Fig. D.1**). While it is true that the models were trained with early stopping (and therefore the training set performance of RCPS could have surpassed that of the post-hoc conjoined models if all models were forced to train for the same number of iterations), we can still draw a contrast between the trend we observed with trained conjoined models, for which we found clear cases where the training-set performance surpassed that of the post-hoc conjoined models even though the test-set performance was worse. Thus, it remains unclear whether the mediocre performance of RCPS on these tasks can be attributed to a tendency to overfit vs. more general optimization difficulties. Further, when we reduced the effective number of filters in the RCPS models to equal the number of filters in standard models, we consistently observed a drop in test-set performance for the RCPS models (**Sec. D.5**), again suggesting that overfitting perse may not be the culprit.

## 5. Discussion

### 5.1. How to extend RCPS to fully-connected layers

When comparing RCPS models to Conjoined models, one apparent point of difference that a reader may notice is that Conjoined models merge the representations of the forward and RC strands after the last fully-connected layer, whereas all existing papers that use RCPS to predict a scalar output perform a representation merging at or before the first dense layer (Shrikumar *et al.*, 2017; Brown and Lunter, 2019; Bartoszewicz *et al.*, 2020). This merging of RCPS representations at the first fully-connected layer is motivated by an implicit assumption that weights at the first fully-connected layer should be roughly symmetric around the positional axis (and therefore it should OK to merge the representations of the forward and RC strands at this layer). However, there may be some cases where the weights of this fully-connected layer are not expected to be symmetric, such as when dealing with regulatory elements that have a strong directional asymmetry (for example, proximal promoter elements that have been oriented such that the transcription start site is on the same strand as the provided input sequence). In such a situation, it may be advantageous to perform representation merging at a later stage. Fortunately, it is easy to extend the RCPS framework to account for fully-connected layers, because such layers can be viewed as a special case of convolutional layers where the receptive field equals the full length of the input. Thus, extending RCPS to full-connected layers simply requires implementing the fully-connected layers as convolutional layers with the appropriate receptive field. Put differently, the “channel” axis of the convolutional layer maps onto the number of units in the fully-connected layer, and the “length” axis of the convolutional layer disappears due to having a size of 1.

### 5.2. A unified description of RC-equivariant models

In this section, we will develop a unified description of reverse-complement architectures present in the literature. Before we do so, it is useful to recap how we generalize the concept of a “reverse-complement” to higher convolutional layers. The key insight introduced in Shrikumar *et al.* (2017) is that if the *c*^*th*^ filter from the beginning of the channel axis can recognize the reverse-complement of the pattern recognized by the *c*^*th*^ filter from the end of the channel axis, then flipping the length and channel axes at an intermediate convolutional layer is equivalent to recomputing the activations of that layer on the reverse-complement input sequence. Note that this definition of a “reverse-complement” encompasses one-hot encoded input sequences when the input is encoded using the ordering ACGT (which is what we use in this work): an A represents the reverse-complement of T, and a C represents the reverse-complement of G. When networks satisfy this property at all convolutional layers, they are said to be “equivariant” under reverse-complementation (Cohen and Welling, 2016; Brown and Lunter, 2019); formally, if we define the revcomp operation to mean “flip the length axis and channel axis”, and we use *f* (*S*) to represent the output of convolutional layer *f* on sequence *S*, then revcomp(*f* (*S*)) = *f* (revcomp(*S*)). RCPS architectures design the convolutional layers to satisfy equivariance by pairing the weights of the forward and “RC” filters using the approach illustrated in **Fig. 2**.

#### 5.2.1. THE CONJOINED-REVCOMP “WRAPPER”

We now introduce the Conjoined-RevComp “Wrapper” (“CJRCWrapper”), which can be viewed as a generalization of Conjoined models that can accept the output of an intermediate model layer as its input, so long as the notion of a reverse-complement is defined for said input. A CJRCWrapper contains two components: a submodel followed by a merge operation. When given an input, the CJRCWrapper proceeds as follows: first, the output of the submodel is computed on the input (call this “orig out”). Then, the reverse-complement of the input is calculated (if the equivariance scheme described above is followed, this is done by flipping both the length and channel axes), and the reverse-complemented input is also provided to the submodel; call the result “rc out”. Finally, “orig out” and “rc out” are combined using the merge operation (ideally in a way that maintains the equivariance). Note that the CJRCWrapper places no constraints on the nature of the submodel; any submodel (even one without convolutional layers) can be used. We now describe how architectures in the literature can be viewed from the perspective of the CJRCWrapper.

#### 5.2.2. RCPS IN THE CJRCWRAPPER PARADIGM

In an RCPS convolution, the *c*^*th*^ filter from the beginning of the convolutional stack is paired with a corresponding “RC” filter at the *c*^*th*^ position from the end of the convolutional stack that can recognize the reverse-complement of what the forward filter recognizes. It follows that the RCPS convolution is a CJRCWrapper around a “submodel” comprised of a single convolutional layer consisting of the forward convolutional filters, where the ‘merge’ operation proceeds as follows: flip the length and channel axes of “rc out”, then concatenate “orig out” with “rc out” along the channel axis. The flipping along the length and channel axes prior to concatenation is sufficient to maintain equivariance in the output. This is illustrated in **Fig. 5**.

**Figure 5.**
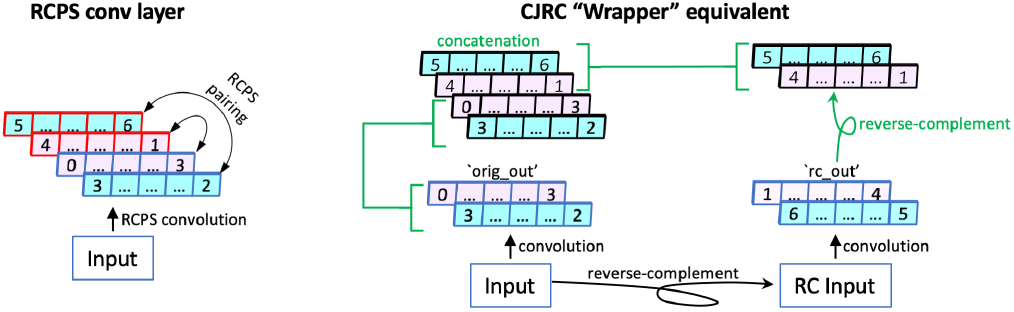
RCPS conv layer implemented in the CJRCWrapper paradigm (Sec. 5.2.1). Cells with similar shading represent filters with shared parameters. Blue outlines denote “forward” filters and red outlines denote paired “RC” filters. Green text denotes the CJR-CWrapper’s “merge” operations. An RCPS conv layer is equivalent to a CJRCWrapper around a “submodel” containing the forward convolutions, where the merge involves reverse-complementing “rc out” and concatenating with “orig out” along the channel axis.

What about the case where RCPS is followed by RC batch normalization (Shrikumar *et al.*, 2017; Bartoszewicz *et al.*, 2020)? The key property of RC batch normalization is that the batch normalization parameters are shared between the forward and RC channels. This can be implemented as a CJRCWrapper around a submodel containing two layers: a single convolutional layer followed by standard batch-normalization. Similarly, RCPS followed by RC dropout (Brown and Lunter, 2019) is equivalent to a CJRCWrapper around a submodel containing a convolutional layer and a dropout layer (with the caveat that the dropout mask is preserved when the submodel is run on both the forward and RC inputs).

RCPS architectures used for binary classification additionally involve a layer that merges the forward and RC representations - for example, Brown and Lunter (2019) had “RC max-pooling”, and in this work we take an elementwise sum over the forward and RC channels. Such layers are equivalent to a CJRCWrapper around a submodel that is just the identity function, followed by a merging op that takes the elementwise maximum (or summation) across “orig_out” and “rc_out”.

FRSS layers (Onimaru *et al.*, 2020) also involve RC parmeter sharing, but are implemented slightly differently, and are equivalent to a CJRCWrapper around two standard convolutional layers where the merge operation is a flipping of “rc_out” along the length axis followed by an elementwise summation. This is discussed more in **Sec. B**.

#### 5.2.3. CONJOINED IN THE CJRCWRAPPER PARADIGM

In contrast to RCPS models, which can be viewed as containing several CJRCWrapper layers that each enclose submodels containing at most one convolutional layer, Conjoined models have a single CJRCWrapper that encloses a sub-model containing all the convolutional layers. In the case of Conjoined models for binary classification tasks, the merge operation for the CJRCWrapper is a simple elementwise operation, such as an average in the case of FactorNet (Quang and Xie, 2019), or maxpooling in the case of DeepBind (Alipanahi *et al.*, 2015). When we extend Conjoined architectures to BPNet-style models, our merge operation is a flipping of the strand and length axes, followed by the elementwise sum.

#### 5.2.4. A SPECTRUM OF ARCHITECTURES BETWEEN RCPS AND CONJOINED

We have shown that each layer in an RCPS models is essentially a CJRCWrapper around a single convolutional layer (optionally followed by batch normalization or dropout), while a Conjoined model is a single CJRCWrapper around an entire submodel that can contain multiple convolutional layers. This leads to a natural intermediary, which is a model with multiple CJRCWrappers that can each contain more than one convolutional layer. If the merging operation between each CJRCWrapper is done in a way that maintains equivariance (e.g. by flipping the length and channel axes of the “rc_out” before concatenating the along the channel axis), the resulting model would also maintain RC equivariance.

Inspired by this observation, we explored the performance of a hybrid CJ-RCPS model on the base-pair-resolution signal profile prediction task. As a reminder, the standard profile prediction models consist of 11 convolutional layers (one standard convolution, followed by 6 dilated convolutions, followed by two convolutional layers). The last two convolutional layers each have two filters (one for the forward strand and one for the RC strand; see **Sec. C.2.2**). In our hybrid model, we enclosed the first seven convolutional layers in a CJRCWrapper, while the last two convolutional layers followed the RCPS formulation (see **Fig. 3**). We observed that this hybrid model tended to outperform trained conjoined models but was not significantly better than full RCPS and remained worse than post-hoc conjoined models (**Fig. D.2**). We defer a more complete exploration of the full space of hybrid architectures to future work.

### 5.3. Conclusion

In this work, we showed that post-hoc conjoined models (trained with RC data augmentation) consistently perform as well as or better than models that were conjoined during training, likely because models that were conjoined during training are more susceptible to overfitting. In fact, post-hoc conjoined models achieved the best or second-best performance across all datasets, surpassed only by RCPS on select datasets. Unfortunately, RCPS was unreliable, in that it sometimes failed to outperform standard models trained with RC data augmentation - particularly for profile prediction. This occasional mediocre performance of RCPS is not due to a representational limitation, given that the RCPS models are capable of representing the solution learned by their conjoined counterparts.

As qualitative support for our observations on the unreliable performance of RCPS, we note Brown and Lunter (2019) used a non-standard initialization of the final output layer to encourage their RCPS models to converge to good solutions. Even so, Brown & Lunter found RCPS did not significantly outperform standard models trained with RC data augmentation on *in vivo* binding data, despite the fact that RCPS did better on simulated data. These apparent difficulties may prevent the full potential of RCPS from being realized out-of-the-box. We thus recommend that deep learning practitioners exercise caution when adopting RCPS into their architectures, and always make sure to compare RCPS against a baseline of post-hoc conjoined models.

We also presented a unified view of conjoined and RCPS architectures, and use it to elucidate a class of architectures that gradually interpolate between fully-conjoined and fully-RCPS models while maintaining RC equivariance. We explored an instantiation of this type of model on the base-pair-resolution profile prediction dataset, and found that while it improved performance relative to trained conjoined models, it did not outperform post-hoc conjoined models. We defer a thorough exploration of this new class of models to future work.

## Author Contributions

HZ performed all model training and evaluation, with mentorship from AS. HZ built the codebase based on code provided by AS. AS conceived of the project, devised the novel RC-equivariant architectures, designed the experiments (with input from HZ), and performed the theoretical analysis. AK provided guidance and feedback. HZ, AS & AK wrote the manuscript.

## Acknowledgments

We thank Anna Shcherbina for assistance with setting up a cloud environment. We thank Alex M. Tseng for providing code to compute performance metrics of profile prediction models. We thank Kelly Cochran for feedback on the manuscript.

## Appendices

### A. RCPS models can be used to represent equivalent conjoined models

In this section, we will show that the RCPS models considered in this work are capable of representing the solutions learned by the corresponding conjoined models that they are benchmarked against. The high-level intuition for our approach is as follows: recall that for each filter in an RCPS model, a corresponding “reverse-complement” filter is created through weight sharing. We will design the weights of our RCPS model such that (1) the activations of the RCPS “forward” convolutional filters on an input *S* match up with the activations of the standard model’s convolutional filters on input *S*, and (2) the activations of the RCPS “reverse-complement” convolutional filters on an input *S* match up with the activations of the standard model’s convolutional filters on *S′* (where *S′* is the reverse-complement of *S*). We will then show how there is a mapping between linear operations in the RCPS model (that occur between the convolutional layers and the final output) and the step where the conjoined model averages the output of the standard model on *S* and *S′*. Thus, the RCPS model can represent any function learned by the corresponding conjoined model.

First, we will recap how RCPS models are constructed. Let **W**^*l,c*^ denote the weights of convolutional channel *c* (0-indexed) in layer *l* of the RCPS model, and let *C*_*l*_ denote the total number of channels (including additional reverse-complement channels generated by RCPS) in layer *l* of the RCPS model. The matrix **W**^*l,c*^ has dimensions of (*w, C*_*l*−1_), where we use *w* to denote the width of the convolutional filters (without loss of generality, we will assume all layers use filters of the same width *w*; we will also set *C*_0_ = 4 to represent the number of channels used in the one-hot encoding of ACGT in the input layer). Under RCPS, the weights of **W**^*l,c*^ are tied to the weights of 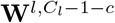. Specifically, if we used 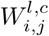 to denote the convolutional kernel weight on position *i* and input channel *j*, and use *b*^*l,c*^ to denote the bias term for channel *c* in layer *l*, then we have:

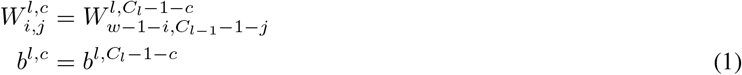

This weight sharing ensures that filter (*l, C*_*l*_−1−*c*) will recognize the reverse-complement of whichever pattern is recognized by filter (*l, c*).

Now, let us extend this notation to the corresponding standard models. Let **W**^\**,l,c*^ denote the weights of convolutional channel *c* at layer *l* of the standard model, and let 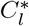 denote the total number of channels in layer *l* of the standard model. In all our benchmarks, we had 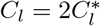 (this was due to the duplication of filters in the RCPS models caused by reverse-complement weight sharing; we also ran comparisons where 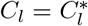 (**Sec. D.5**) - however, when 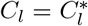, the equivalence explained in this section does not hold). Let us further use 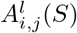 and 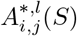 to denote the activations in layer *l*, position *i*, channel *j* for the RCPS and standard model respectively when sequence *S* is supplied as input. When *l* = 0 (denoting the input layer), we will pretend 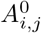 and *A*^*,0^ are simply the identity function. Let us also use *S′* to denote the reverse-complement of *S*, and let *L*_*l*_ denote the length of layer *l*.

If we set the weights for channels *c* = 0 through 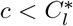 in the convolutional layers of the RCPS model such that:

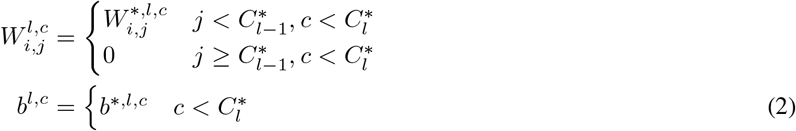

And if we also set the weights for channels 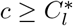 through *c < C*_*l*_ in accordance with RCPS weight sharing (**Eqn. 1**) such that:

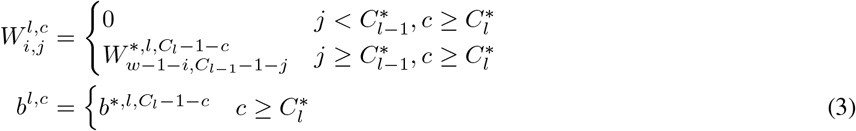

Then we can prove that:

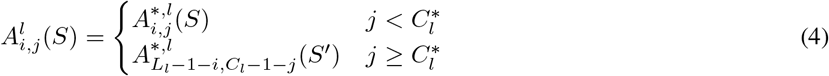

#### A.1. Proof of Eqn. 4

We will prove this by induction. We note that **Eqn. 4** holds true in the case of the input layer *l* = 0, assuming that one-hot encoding was done using the ordering ACGT. As mentioned, *A*^0^ and *A*^*,0^ are simply the identity function, so 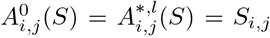. **Eqn. 4** simply states that if there is an A(*j* = 0) or a C(*j* = 1) at position *i* in the input sequence *S*, there will respectively be a T(*C*_*l*_ −*j* −1 = 4−1 = 3) or a G(*C*_*l*_ −*j*−1 = 4−2 = 2) at position *L*_0_−1−*i* of the reverse-complement *S′*. Here *L*_0_ is simply the length of the input sequence. Thus, **Eqn. 4** holds for the base-case of *l* = 0 due to the reverse-complement property of DNA.

We will now show that if **Eqn. 4** holds for the base-case of *l* = 0, it holds for all *l >* 0. From the definition of a convolutional operation, we have:

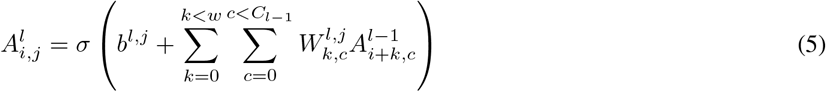

Where *σ* denotes a nonlinearity. Let us begin by proving the case where *j* (the index of the convolutional channel) satisfies 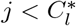 (i.e. *j* is in the first half of the convolutional filters - recall that 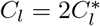). From **Eqn. 2**, we have:

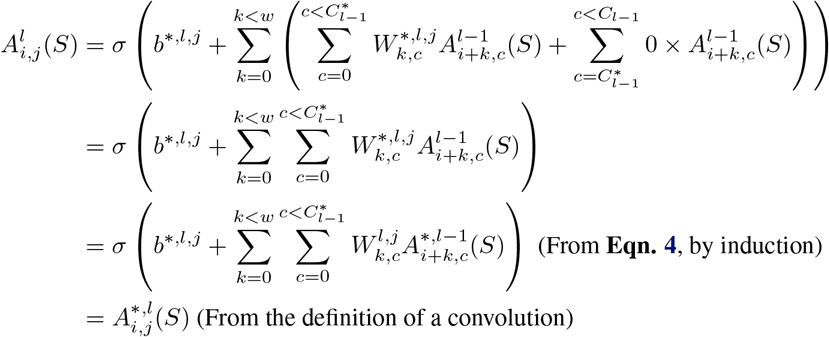

Let us now prove the case where 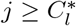. Substituting **Eqn. 3**, we have:

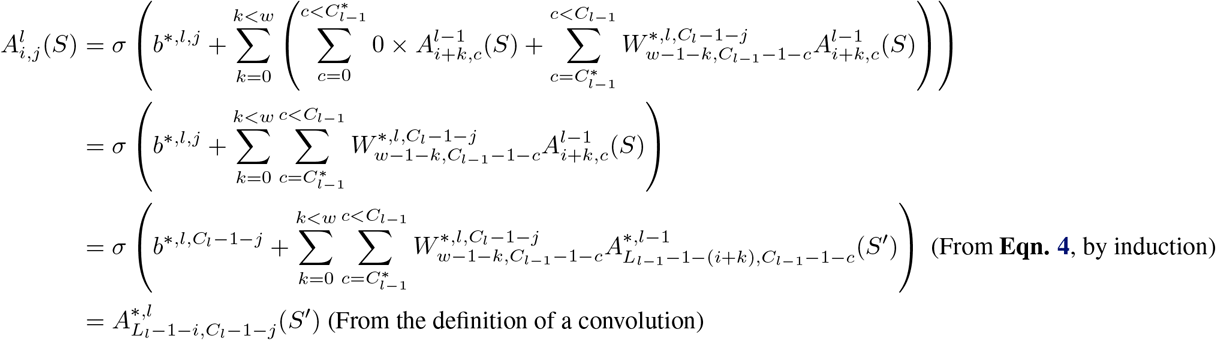

Thus, we have proven **Eqn. 4** for any layer in a stack of convolutions, extending from the input upwards. In words, this shows that the convolutional activations of the standard model on the forward sequence *S* correspond to the convolutional activations of the “forward” filters of the RCPS model on the sequence *S*, and that the convolutional activations of the standard model on the RC sequence *S′* correspond to the convolutional activations of the “RC” filters of the RCPS model on sequence *S*.

#### A.2. Combining the representations on the forward and reverse strands

Let us now consider how the conjoined model combines the representations on the forward and reverse strands. In binary conjoined models, the stack of convolutional layers is followed by a linear transformation that predicts the logit of the sigmoid, after which the representations from both strands are averaged and passed through the sigmoid. Specifically, if we use 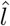 to denote the last convolutional layer, *g* to denote the function computing the logit in the standard model, and we use **W**^*,**g**^ & *b*^\**,g*^ to denote the weights & biases of *g*, we have:

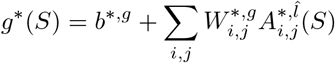

Thus, the corresponding output *g*^**^(*S*) of the conjoined model is:

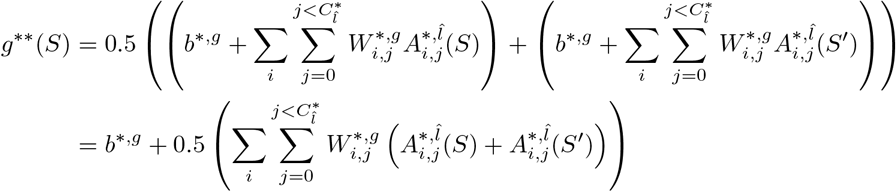

In the case of the RCPS binary models, the “forward” and “RC” channels at the end of the stack of convolutional layers are added together (after reverse-complementing the RC channels to be compatible with the forward channels - see **Fig. 1**), and then a linear operation is applied to obtain the logit of the sigmoid. If we let *g* denote the function computing the logit of the RCPS model, and use **W**^**g**^ & *b*^*g*^ to denote the weights and biases of *g*, we have:

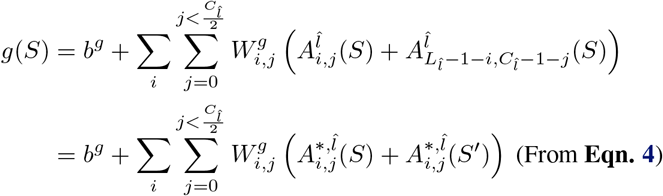

The reason we iterate from *j* = 0 to 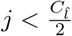 is that the effective number of filters in the RCPS model gets halved when the forward and reverse-complement channels are added together. Also recall that 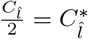. If we therefore set *b*^*g*^ = *b*^\**,g*^ and 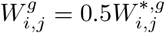, we will achieve *g*(*S*) = *g*^**^(*S*).

In the case of the profile prediction models, there is an additional nuance that separate predictions are made for the two output strands. For simplicity, we will treat the control bias track as though it is another channel at the end of the last nonlinear convolutional layer. We will denote the activations of the last nonlinear convolutional layer in the standard profile prediction model using 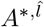. Since the operations following the last nonlinear convolutional layer are all linear convolutions (see **Sec. C.2.2**), we will represent the output of the strands at position *i* for the standard model as:

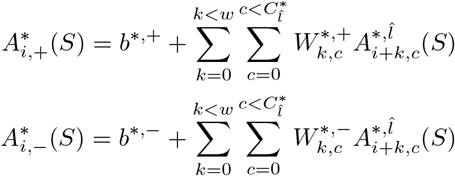

After the reverse strand predictions are flipped and averaged with the forward strand, the output of the strands for the conjoined model is:

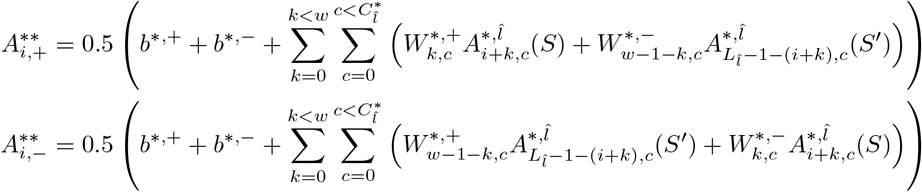

By comparison, the output of the forward strand for the RCPS model is written as:

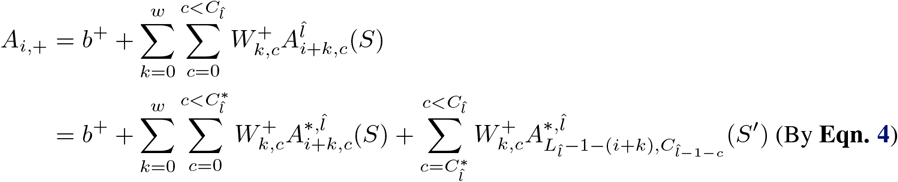

If we thus set the weights such that *b*^+^ = 0.5(*b*^*,+^ + *b*^*,−^) and

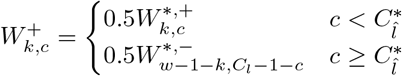

We achieve 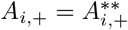. The weights for *A*_*i*,−_ would be given by **Eqn. 2**, and a similar calculation can be done to show 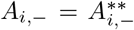. Thus, the RCPS models can be used to represent the functions learned by the corresponding conjoined models.

### B. FRSS layers in the CJRCWrapper paradigm

Onimaru *et al.* (2020) proposed FRSS (Forward and Reverse Sequence Scan) layers as a form of RC parameter sharing. In FRSS layers, a one-hot encoded input sequence is supplied to two “branches”. The first branch applies two standard convolutional layers (including any associated pooling and nonlinearities) to the input. The second branch is similar to the first branch, but runs the convolutional operations with “rotated” kernel weights, such that the filters in the second branch recognize the reverse-complement of whatever patterns their corresponding filters in the first branch recognize. The outputs of the two branches are then combined via an elementwise summation before being supplied to the rest of the network layers (which may include additional pooling, convolutional and fully-connected operations).

The FRSS layer is equivalent to a CJRCWrapper around two standard convolutional layers, where the merge operation is a flipping of “rc out” along the length axis (to positionally align the forward and RC representations), followed by an elementwise summation. Although it may seem intuitive to positionally align the forward and RC representations prior to summing them, doing so does not guarantee RC equivariance later in the model. To appreciate why, let us presume that the forward version of the input sequence contains a motif at a location that is *x*bp from the start of the sequence; this means the RC version of the input sequence will contain a motif at *x*bp from the end of the sequence. Now consider what happens when the representation learned by the FRSS layer is eventually supplied to a fully-connected layer: unless the fully-connected layer learns to place equal weight on both locations in the input sequence, the activations of the fully-connected layer may differ between the forward and RC strands. In practice, fully-connected layers have been observed to place asymmetrical importance on positions that are equidistant from the center of the input sequence, even when those positions would be given equal weight by an ideal model (Alexandari *et al.*, 2017). To avoid this issue and guarantee RC equivariance, our RCPS models flip the positional axis of the RC filter activations prior to merging the forward and RC channels (similar to what was done in Brown and Lunter (2019)). Doing so breaks the positional alignment of the forward and RC filters, which is why the merging of forward and RC filters is RCPS models is performed only immediately prior to the first fully-connected layer (it is presumed that the fully-connected layers mainly focus on the locations of convolutional filter activations relative to the center of the sequence, which is still preserved by the flipping).

### C. Supplementary Methods

#### C.1. Binary prediction

##### C.1.1. BINARY PREDICTION DATASETS

Our simulated datasets were based on those used in Shrikumar *et al.* (2017), but with a few slight modifications. Three “sets” of sequences were generated using the simdna package (Shrikumar *et al.*, 2019); one contained instances of the GATA known6 Position Weight Matrix (PWM), one contained instances of the ELF1 known2 PWM, and one contained instances of RXRA known1 (motifs were taken from Kheradpour and Kellis (2014)). 10,000 sequences were generated for each set, for a total of 30,000 sequences. The sequences were simulated as follows: (1) generate a random background sequence of length Xbp (where X was either 200 or 100) with 40% GC content (2) determine the number of instances of the motif to insert by sampling from a truncated Poisson distribution with mean 2, max 3 and min 1 (3) sample the motif instances from the specified PWM (either GATA known6, RXRA known1 or ELF1 known1), (4) reverse complement each sampled instance with probability 0.5 (5) insert each sampled instance at a random position within the sequence, with the constraint that it does not overlap a previously inserted instance. We randomly allocated 30% of the data to validation, 30% to testing, and 40% to training. Labels where then generated as follows: there were three binary tasks, each task corresponding to a particular set; a sequence was labeled as a 1 for a task if it originated from the corresponding set, and with a 0 otherwise.Finally, we simulated mislabeling noise by flipping each individual label with 20% probability (this approach of adding noise differed slightly from the approach used in (Shrikumar *et al.*, 2017) in that, in our case, the probability of flipping the label for a particular task was independent of the probability of flipping the label for the other tasks).

Our processed TF ChIP-seq datasets were identical to those created by Shrikumar *et al.* (2017), where the raw data was produced by the ENCODE consortium (ENCODE Project Consortium, 2012). For Ctcf, the file used was “wgEncodeBroad-HistoneGm12878CtcfStdAlnRep0”, for Spi1, the file used was “wgEncodeHaibTfbsGm12878Pu1Pcr1xAlnRep0”, and for Max, we used “wgEncodeSydhTfbsGm12878MaxIggmusAlnRep0”. The positive and negative sets were prepared as follows: for the positive set, we used 1000bp windows centered around the summits of rank-reproducible peaks (Landt *et al.*, 2012). For the negative set, we used 1000bp around the summits of DNase peaks in Gm12878 that did not overlap the top 150K relaxed ChIP-seq peaks of the TF (the “relaxed” peaks were called by SPP at a 90% FDR). The file used for DNase peaks was “E116-DNase.macs2.narrowPeak.gz”, produced by the Roadmap consortium (Roadmap Epigenomics Consortium *et al.*, 2015). The training set consisted of all chromosomes except chr1 & chr2, the validation set consisted of chr1, and the testing set consisted of chr2. The Max dataset had 12,542 positives and 206,628 negatives, the CTCF dataset had 44,982 positives and 225,533 negatives, and the SPI1 dataset had 42,938 positives and 203,960 negatives.

##### C.1.2. BINARY PREDICTION MODELS

Our model architectures for the binary datasets were based on the ones from Shrikumar *et al.* (2017). All standard models trained on the simulated data employed one convolutional layer with 20 filters of kernel width 21 and stride 1, followed by batch normalization and the ReLU nonlinearity, followed by maxpooling with width and stride 20, followed by the sigmoid output layer with 3 neurons. All standard models trained on the TF ChIP-seq data had three 16-filter stride-1 convolutional layers of kernel widths 15, 14 & 14 respectively, each of which was accompanied by batch normalization and a ReLU nonlinearity, followed by maxpooling of width 40 & stride 20, followed by the single sigmoid output.

For the conjoined models trained on binary prediction tasks, we averaged the predicted sigmoid logits across both forward and RC inputs prior to passing the result through a sigmoid function to obtain the final prediction. Note that this differs slightly from prior work (Alipanahi *et al.*, 2015; Quang and Xie, 2019); in Alipanahi *et al.* (2015), the maximum prediction is taken across strands, while in Quang and Xie (2019), predictions are averaged after the sigmoid nonlinearity is applied. We justify taking the average rather than the maximum as this was found to produce superior results both in Bartoszewicz *et al.* (2020) and in our own benchmarking (**Sec. D.6**); we suspect this is because averaging results in more informative gradient updates for conjoined models during training (if the “max” is applied across both strands, then during training, the gradient of the model with respect to one of the strands would always be zero). We justified taking the average of the sigmoid logits rather than taking the average of the post-sigmoid probabilities because of the equivalence proven in **Sec. A**, and also because of the following probabilistic interpretation: the logit of a sigmoid represents the log of the predicted odds ratio that the input belongs to the positive class; averaging sigmoid logits can thus be interpreted as taking the geometric mean of two predicted odds ratios. In our benchmarking, we found that taking the average of the logits performed comparably to (if not slightly better than) taking the average of the post-sigmoid probabilities (**Sec. D.6**).

For the RCPS architectures trained on binary prediction tasks, we summed representations across strands prior to the final sigmoid output layer (**Fig. 1**). This is similar to Bartoszewicz *et al.* (2020) and is functionally equivalent to the “weighted sum” layer used in Shrikumar *et al.* (2017) to merge representations, but with a more simplified implementation. Because taking the sum rather than the average can impact the learning rate dynamics, we verified that using the average of the representations did not produce substantially different results than using the sum (**Sec. D.6**).

##### C.1.3. BINARY PREDICTION TRAINING HYPERPARAMETERS

Following Shrikumar *et al.* (2017), all binary prediction models were trained with a binary cross-entropy loss and the Adam optimizer with the default Keras learning rate of 0.001. In the case of the simulated binary datasets, we replicated the setup from Shrikumar *et al.* (2017) and defined our “epochs” to be 5000 training examples (note: canonically, an “epoch” is defined to be a single pass through the entire training set - however, when the data is loaded on-the-fly using a generator in order to avoid loading all the data at once, epochs can instead be defined by the number of training examples seen). At a batch size of 500 for the simulated data, each “epoch” therefore corresponded to 10 training iterations. The training was terminated when the validation set loss failed to show improvement over 10 consecutive “epochs”, and the model weights at the “epoch” with the best validation set loss were used for performance comparisons. In the case of the TF ChIP-seq datasets, Shrikumar *et al.* (2017) defined an “epoch” as 5000 training examples, and used batch sizes of 100 - thus, each “epoch” corresponded to 50 training iterations. Shrikumar *et al.* (2017) trained each of the TF ChIP-seq models for 4000 training iterations and used the model weights from the epoch that achieved the best validation set AuROC. *Shrikumar et al.* (2017) also upweighted their positive examples according to the class imbalance (16.47:1 for Max, 4.75:1 for Spi1 and 5.01:1 for Ctcf). We replicated this exact setup, but also explored how changes in these training hyperparameters impacted performance (**Fig. D.3, D.4, D.5** and **Sec. D.3**) - in particular, we set the maximum number of training iterations to be 8000 (in addition to 4000), we selected the best validation set epoch according to prediction loss (in addition to AuROC), and we handled the class imbalance by up**sampling** positive examples to achieve a 1:4 class ratio (rather than up**weighting** positive examples in the loss).

#### C.2. base-pair-resolution signal profile prediction

##### C.2.1. PROFILE PREDICTION DATASETS

We trained our BPNet models using the processed datasets from Avsec *et al.* (2020). These datasets were ChIP-nexus profiles of the pluripotent TFs Sox2, Oct4, Klf4, and Nanog in mouse embryonic stem cells. At each position and strand in each of the four TFs, the profile included the 5’ end read counts. 1000bp windows were selected around the peak summits from the Irreproducible Discovery Rate (IDR) optimal peaks sets (Landt *et al.*, 2012). These sequences were inputted into the BPNet Architecture. Because ChIP-nexus experiments can have certain biases, experimental control data was used from PAtCh-CAP59 (Terooatea *et al.*, 2016). The validation set consisted of regions on chr1, chr8 and chr9, the testing set of regions on chr2, chr3 and chr4, and the training set consisted of all other regions. The dataset sizes for the training, validation and test sets were, 6748, 2084 & 2167 for Sox2, 15946, 4818 & 5085 for Oct4, 35009, 10542 & 10908 for Nanog, and 36201, 10283 & 11117 for Klf4.

##### C.2.2. PROFILE PREDICTION MODELS

BPNet models predict the shape of the observed TF binding signal at base-pair-resolution resolution in a 1kb interval using both DNA sequence and the “control” signal track as inputs. The predicted profiles take the form of a probability distribution over each position, for each strand. This probability distribution is compared to the observed distribution of raw read counts in the interval and is scored using a multinomial loss function.

The original BPNet architecture contained two output heads, where one head predicted the shape of the signal profile at base-pair-resolution resolution, and the other head predicted the total read counts in a given region. For the purpose of controlled benchmarking, we trained models on only the profile prediction head (thereby avoiding differences in performance caused by competition between the two heads). Our BPNet architecture consists of a one-hot encoded input sequence that is supplied to a convolutional layer with 64 filters and a kernel width of 21, followed by a stack of 6 dilated convolutional layers with 32 filters each and a kernel width of 3. The dilation rate of the convolutional layers increased by a factor of 2 with each layer - i.e. the dilations were 2,4,8,16,32 and 64. The output of each layer was also added to the output of the previous layer in order to create the input to subsequent layers. At the end of the stack of dilated convolutional layers (all of which had ReLU activations), a linear convolutional layer with 2 filters and a filter width of 75 was applied (note that this is equivalent to the “deconvolutional” layer used in the BPNet paper; when the stride is 1, as is the case here, a “deconvolutional” layer is equivalent to a convolutional layer). The output of this layer was then concatenated to the control track’s signal profile, and then a second linear convolutional layer with 2 filters and a kernel with of 1 was used to obtain the final profile prediction (one filter each for the positive and negative strands). Note that, from the perspective of the multinomial loss function, these predictions represent the logits of a distribution to which a softmax is implicitly applied (the softmax is applied separately for each strand). All convolutional layers except for the final layer were followed by ReLU nonlinearities. All models were trained using the Adam optimizer and the default Keras learning rate of 0.001 and a batch size of 64. An “epoch” was defined as a full pass through the training dataset, and models were trained using early stopping with a patience of 10 epochs. The metric used for early stopping was the prediction loss on the validation set, and “restore best weights” was set to True (meaning that the model weights at the epoch with the best validation set loss were used). Our handling of reverse-complement equivariance for BPNet-style models is described in **Fig. 3**.

One modification that we made to the BPNet architecture was to avoid zero-padding; in the original BPNet architecture, the inputs to convolutional layers were zero-padded such that the output of the convolutional layer had the same dimensions as its input (this is called “same” padding in keras). The “same” padding was necessary for the residual connections, which perform elementwise additions of different convolutional layers. To avoid zero-padding, we instead supplied a longer initial input sequence (1346bp); in order to make different convolutional layers have compatible dimensions for the residual connections, we trimmed away the ends of the longer layer prior to performing the elementwise addition.

##### C.2.3. METRICS TO EVALUATE PROFILE PREDICTION

Profile model predictions were evaluated according to three metrics: Spearman correlation, Pearson correlation, and Jensen-Shannon divergence. Recall that BPNet’s predicted base-resolution signal profiles take the form of a probability distribution. This predicted distribution is compared against the observed distribution of reads. In the case of Spearman and Pearson correlation, we bin both the true distribution and the prediction distribution into bins of size 1bp, 5bp and 10bp, and compute the correlation for each example, strand and binning resolution (here, “binning” means taking the maximum value of the constitutive elements in the bin). In the case of Jensen-Shannon divergence (JSD), the predicted and true probability distributions are smoothed using a Gaussian kernel with *σ* = 3, and the JSD is computed separately for each example and strand. The reported values for the Spearman correlation, Pearson correlation and JSD are the average of the correlations across all examples, strands and binning resolutions (if applicable).

### D. Supplementary Results

#### D.1. Comparison of Training vs. Test-set Performance for CJ and RCPS architectures

**Figure D.1.**
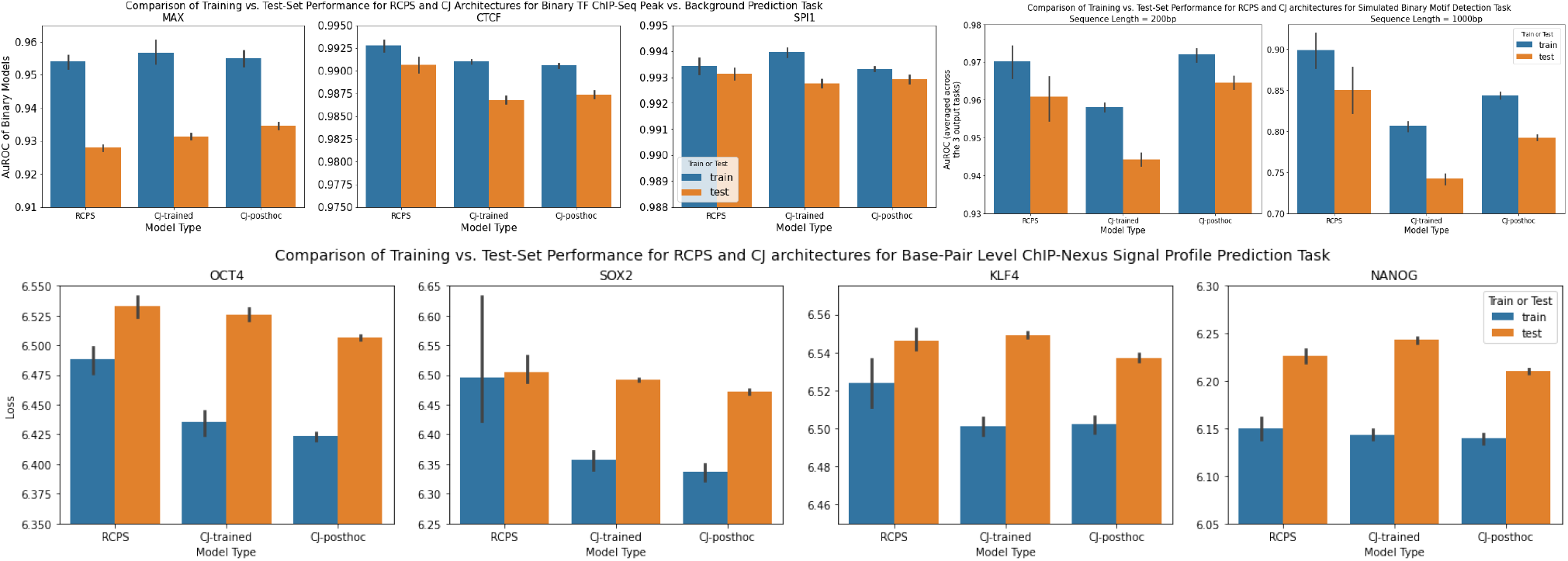
Comparison of training vs. test-set performance for RCPS and CJ architectures. Bar heights represent the average performance over 10 random seeds, and error bars represent the 95% confidence intervals for the mean generated by seaborn (Waskom *et al.*, 2017) using 1000 bootstrapped samples. RCPS denotes RC Parameter Sharing. CJ-trained are models that were conjoined during both training and test time. CJ-posthoc are standard models trained with RC data augmentation that were converted to conjoined models only after training. We show AuROC (rather than AuPRC) for the binary models because AuROC was the criterion used to select the training iteration with the best validation set performance (this is consistent with the training procedure used in Shrikumar *et al.* (2017)). Similarly, for the profile models, we show the crossentropy loss of predicting the positions of the reads (this is equal to the multinomial loss - i.e. the training loss - minus a constant factor; the multinomial loss was used for selecting the best training iteration; lower is better). On the binary TF ChIP-seq datasets and base-pair resolution signal profile prediction models, we see that CJ-posthoc models consistently show a mean test-set performance that is comparable to or better than the mean test-set performance of CJ-trained, coupled with a mean training-set performance that is comparable to or worse than the mean training-set performance of CJ-trained, which matches the hypothesis that CJ-trained has a tendency to overfit to the training set relative to CJ-posthoc. Although the trend is not observed on the simulated data, this could be because the models were trained with early stopping (thus, it is possible that the training set performance of the trained conjoined models may have surpassed that of the post-hoc conjoined models if all models were forced to keep training for a fixed number of iterations). By contrast, on tasks where the mean test-set performance of RCPS comparable to or worse than the mean test-set performance of CJ-posthoc (i.e. all tasks except binary CTCF ChIP-seq, binary SPI1 ChIP-seq, and the simulated data with sequence length 1000bp), we observe that the training set performance of RCPS is never significantly better than that of CJ-posthoc, and (as in the case of the Oct4, Sox2 and Klf4 profile prediction tasks) is sometimes significantly worse.

#### D.2. Performance of a hybrid Conjoined/RCPS architecture on base-pair-resolution signal profile prediction

**Figure D.2.**
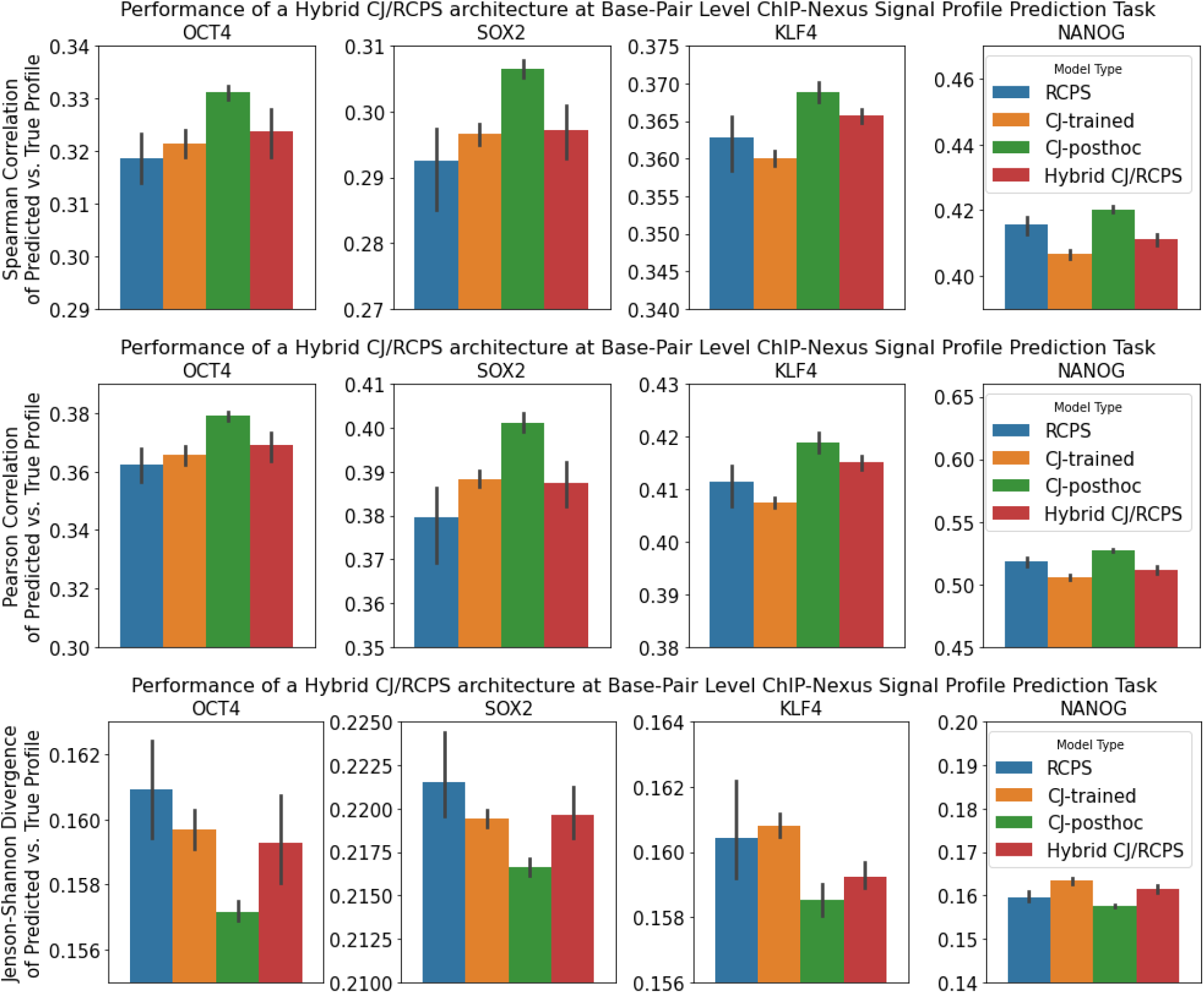
Performance of a hybrid Conjoined/RCPS architecture on base-pair-resolution signal profile prediction. Bar heights represent the average performance over 10 random seeds, and error bars represent the 95% confidence intervals for the mean generated by seaborn (Waskom *et al.*, 2017) using 1000 bootstrapped samples. RCPS denotes RC Parameter Sharing. CJ-trained are models that were conjoined during both training and test time. CJ-posthoc are standard models trained with RC data augmentation that were converted to conjoined models only after training. “Hybrid CJ/RCPS” is the hybrid Conjoined/RCPS architecture described in **Sec. 5.2.4**. Note that, for Jensen-Shannon divergence, lower is better. We observe that this hybrid architecture performs comparably to or better than CJ-trained, but does not significantly outperform the full RCPS model and consistently underperforms relatived to CJ-posthoc.

#### D.3. Some datasets show sensitivity to training hyperparameters

Previous analyses Shrikumar *et al.* (2017) had shown that RCPS architectures outperformed standard architectures with RC data augmentation on several binary output TF binding prediction tasks. However, we found that increasing the maximum number of training iterations (i.e. batch updates) from 4000 (used by Shrikumar *et al.* (2017)) to 8000, resulted in the standard architectures with RC data augmentation outperforming the RCPS architecture for predicting binary binding of the Max protein (**Fig. D.3**). This trend was not observed for Ctcf (**Fig. D.4**) and Spi1 proteins (**Fig. D.5**), where RCPS remained in the lead irrespective of the limit on the number of training iterations. The Max dataset differs most dramatically from Spi1 and Ctcf in that it is has a more acute class imbalance (Max has 16.47:1 positive to negative example ratio compared to 4.75:1 for Spi1 and 5.01:1 for Ctcf). We hypothesize that due to the smaller number of positive training examples, the Max task is inherently harder to learn on, which may explain why the standard architecture needs more training iterations to find a good solution.

Inspired by this result, we explored the effect of other hyperparameters that impact model training (**Fig. D.3, D.5, D.4**). For example, we explored the impact of the metric used to select the “best” validation set epoch. By default, the Keras “early stopping” callback selects the epoch with the best validation set loss. We found that this default approach could occasionally yield significantly lower validation-set auROCs and auPRCs relative to using the epoch with the best validation-set auROC, particularly for the Max dataset. Note that the approach of selecting the best validation set epoch using auROC was used in Shrikumar *et al.* (2017), and we found it was necessary to replicate their results. We also found that when the training-set class imbalance was handled by up**sampling** positive examples to achieve a positives:negatives ratio of 1:4 (instead of up**weighting** positive examples in the loss function according to the class ratios, as was done in Shrikumar *et al.* (2017)), the model auPRCs generally improved for the all three tasks. However, regardless of these hyperparameter choices, the main trends described in this work were robust to these differences in hyperparameters.

**Figure D.3.**
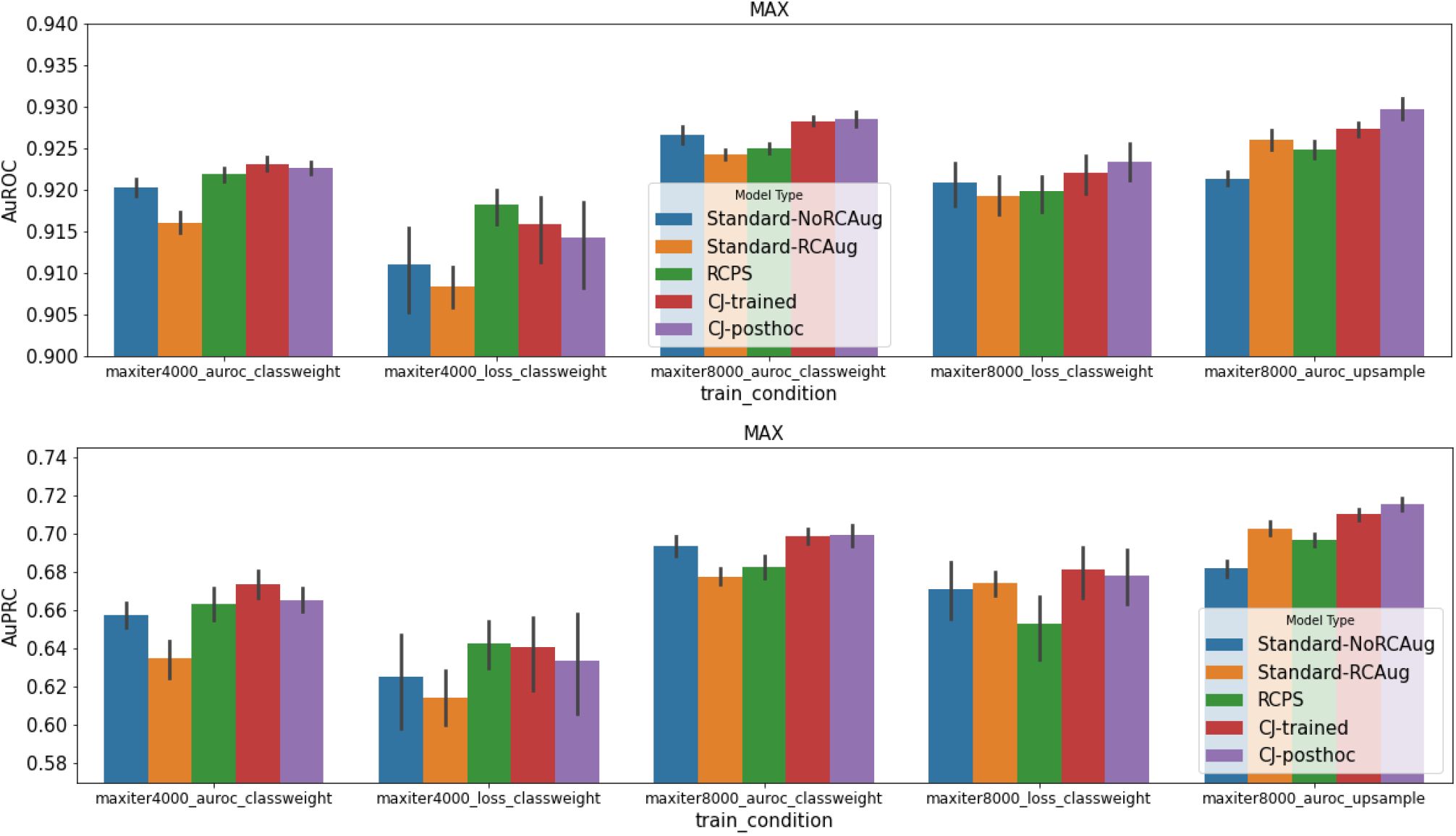
Benchmarking effect of different training hyperparameters for Max TF ChIP-seq binary peak prediction. Bar heights represent the average performance over 10 random seeds, and error bars represent the 95% confidence intervals for the mean generated by seaborn (Waskom *et al.*, 2017) using 1000 bootstrapped samples. “Standard-RCAug” and “Standard-noRCAug” are standard models trained with and without RC data augmentation. RCPS denotes RC Parameter Sharing. CJ-trained are models that were conjoined during both training and test time. CJ-posthoc are standard models trained with RC data augmentation that were converted to conjoined models only after training.

**Figure D.4.**
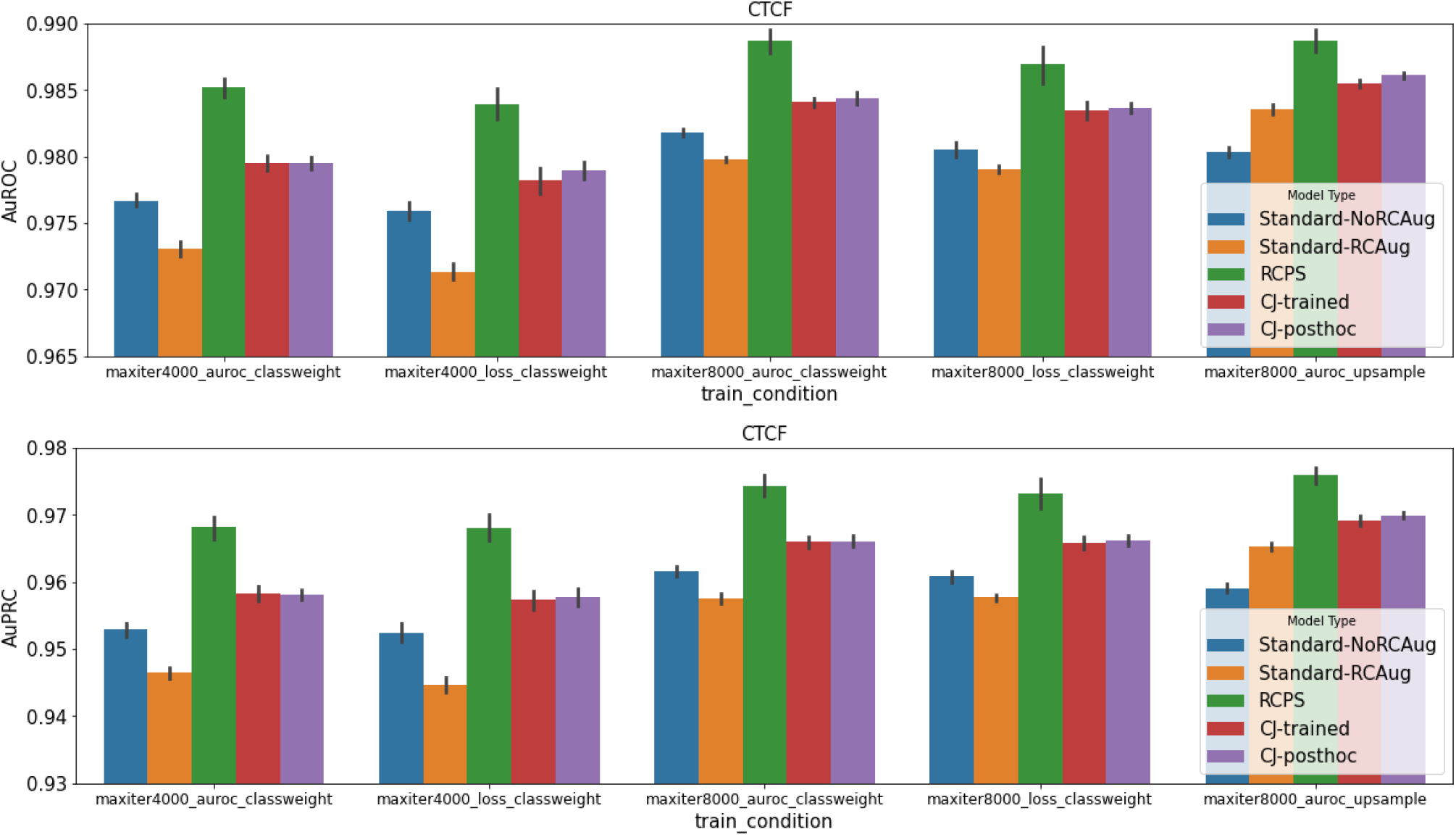
Benchmarking effect of different training hyperparameters for Ctcf TF ChIP-seq binary peak prediction. Bar heights represent the average performance over 10 random seeds, and error bars represent the 95% confidence intervals for the mean generated by seaborn (Waskom *et al.*, 2017) using 1000 bootstrapped samples. “Standard-RCAug” and “Standard-noRCAug” are standard models trained with and without RC data augmentation. RCPS denotes RC Parameter Sharing. CJ-trained are models that were conjoined during both training and test time. CJ-posthoc are standard models trained with RC data augmentation that were converted to conjoined models only after training.

**Figure D.5.**
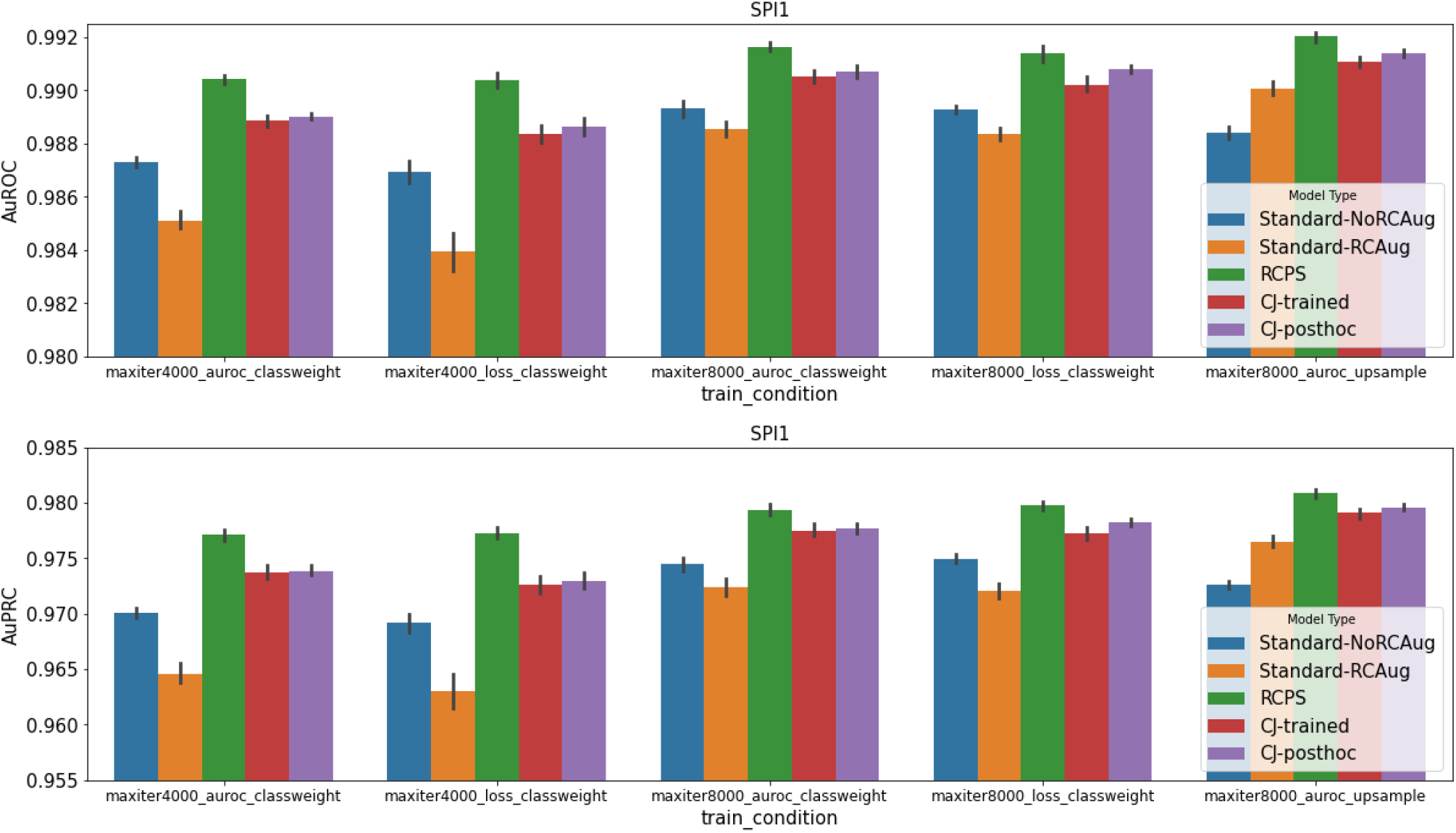
Benchmarking effect of different training hyperparameters for Spi1 TF ChIP-seq binary peak prediction. Bar heights represent the average performance over 10 random seeds, and error bars represent the 95% confidence intervals for the mean generated by seaborn (Waskom *et al.*, 2017) using 1000 bootstrapped samples. “Standard-RCAug” and “Standard-noRCAug” are standard models trained with and without RC data augmentation. RCPS denotes RC Parameter Sharing. CJ-trained are models that were conjoined during both training and test time. CJ-posthoc are standard models trained with RC data augmentation that were converted to conjoined models only after training.

#### D.4. Alternative Performance Metrics for Signal Profile Prediction

Included here are the results of the performance evaluation for signal profile prediction using Pearson correlation and Jensen-Shannon divergence as the metrics. Refer to **Sec. C.2.3** for an explanation of how the metrics are calculated.

**Figure D.6.**
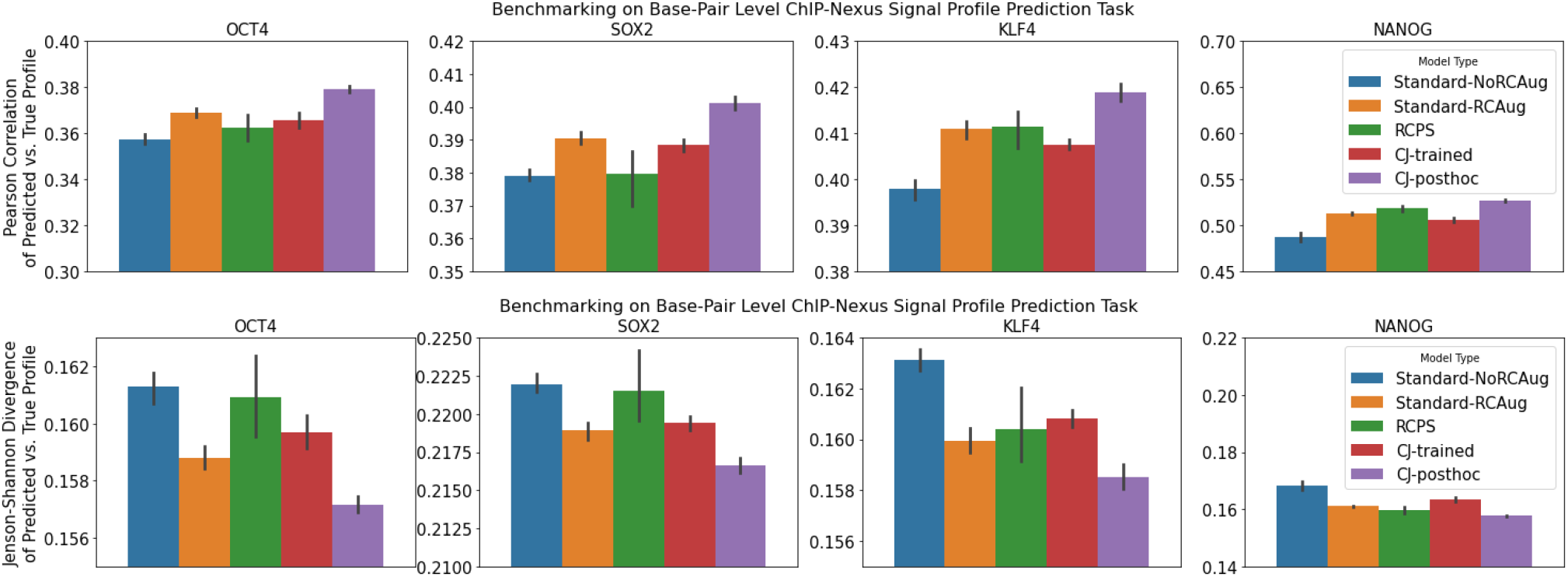
Benchmarking of profile prediction models - Pearson correlation and Jensen-Shannon divergence. Note that, for Jensen-Shannon divergence, lower is better. Results with Spearman correlation as the metric are in **Fig. 4**. The metrics are explained in **Sec. C.2.3**. Bar heights represent the average performance over 10 random seeds, and error bars represent the 95% confidence intervals for the mean generated by seaborn (Waskom *et al.*, 2017) using 1000 bootstrapped samples. “Standard-RCAug” and “Standard-noRCAug” are standard models trained with and without RC data augmentation. RCPS denotes RC Parameter Sharing. CJ-trained are models that were conjoined during both training and test time. CJ-posthoc are standard models trained with RC data augmentation that were converted to conjoined models only after training.

#### D.5. Performance benchmarks of RCPS with a reduced number of filters

As discussed in **Sec. 3.1**, in the case of the RCPS architectures, each filter has a reverse-complement counterpart that is created at runtime (via weight sharing), which increases the representational capacity of the model. Thus, the “effective” number of filters of the RCPS model could be considered to be twice that of the standard models. To account for this, we also trained RCPS models with half the number of filters (such that the “effective” number of filters would be comparable to that in standard models). However, on all tasks we evaluated, we consistently observed that this decreased performance (**Fig. D.7**, **Fig. D.8**).

For the TF ChIP-seq binary datasets, this difference was most apparent in the Max TF ChIP-seq binary task, where the reduction in filters caused the RCPS models with half the number of filters to perform considerably worse than all other models, including standard models trained without data augmentation. This trend is consistent with the generally poor performance of RCPS on the Max TF ChIP-seq dataset (**Sec. D.3**). For the Ctcf ChIP-seq binary dataset, the performance of RCPS models with half the number of filters was still superior to the other non-RCPS models benchmarked, and for Spi1 ChIP-seq the RCPS models with half filters retained comparable performance to the best non-RCPS models.

**Figure D.7.**
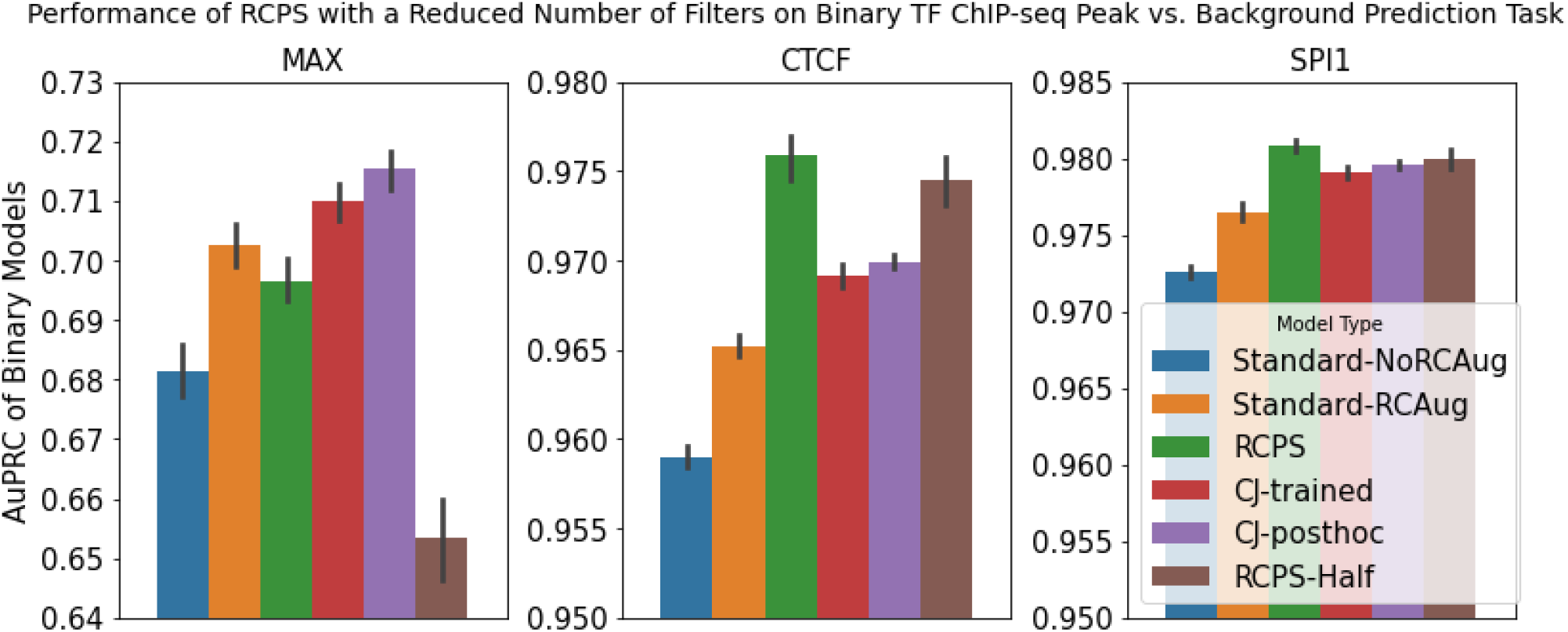
Performance of RCPS models with half the number of filters on binary TF ChIP-Seq datasets. Bar heights represent the average performance over 10 random seeds, and error bars represent the 95% confidence intervals for the mean generated by seaborn (Waskom *et al.*, 2017) using 1000 bootstrapped samples. “Standard-RCAug” and “Standard-noRCAug” are standard models trained with and without RC data augmentation. RCPS denotes RC Parameter Sharing. CJ-trained are models that were conjoined during both training and test time. CJ-posthoc are standard models trained with RC data augmentation that were converted to conjoined models only after training. RCPS-half are models with the same architecture as regular RCPS models but with half the number of filters. Similar to **Fig. 4**, training hyperparameters were set to the tested combination that tended to produced the highest overall AuPRCs.

**Figure D.8.**
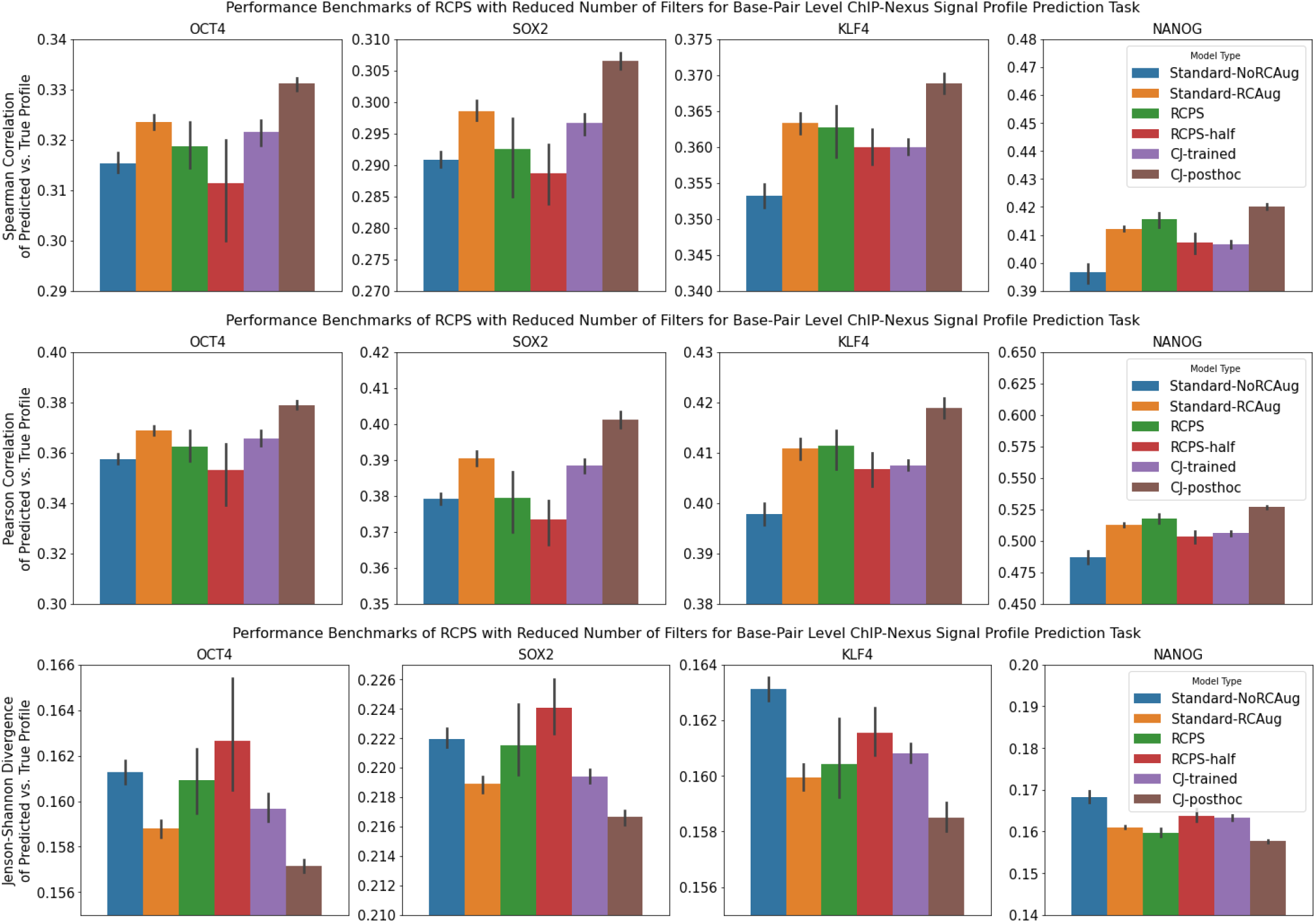
Performance of RCPS models with half the number of filters on profile prediction datasets. Bar heights represent the average performance over 10 random seeds, and error bars represent the 95% confidence intervals for the mean generated by seaborn (Waskom *et al.*, 2017) using 1000 bootstrapped samples. “Standard-RCAug” and “Standard-noRCAug” are standard models trained with and without RC data augmentation. RCPS denotes RC Parameter Sharing. CJ-trained are models that were conjoined during both training and test time. CJ-posthoc are standard models trained with RC data augmentation that were converted to conjoined models only after training. RCPS-half are models with the same architecture as regular RCPS models but with half the number of filters.

#### D.6. Comparison of Different Representation Merging Strategies for Binary Models

For thoroughness, we compared the performance of conjoined and RCPS models using different strategies for combining the predictions of the two branches. In addition to standard models trained with and without data augmentation, we looked at the following cases:

- RCPS models where the representations of the forward and RC strands were combined via summation (as was done in this work). This was denoted as “RCPS” in the figure legend.
- RCPS models where the representations of the forward and RC strands were combined via averaging (this is representationally equivalent to summing, but we wanted to test if it impacted learning dynamics due to scaling of the learning rate). This was denoted as “RCPS-Avg”.
- Trained conjoined models where the predictions on the forward and RC strands were averaged at the level of the sigmoid logit (as was done in the remainder of this work), prior to applying the final sigmoid nonlinearity. This was denoteed as “CJ-trained”.
- Trained conjoined models where the maximum prediction across the forward and RC strand was taken (rather than combining the strand predictions by averaging the logits), as was done in Alipanahi *et al.* (2015). This was denoted as “CJ-trained-max”.
- Post-hoc conjoined models where the predictions on the forward and RC strands were averaged at the level of the sigmoid logit (as was done in the remainder of this work), prior to applying the final sigmoid nonlinearity. This was denoted as “CJ-posthoc”.
- Post-hoc conjoined models where the predictions on the forward and RC strands were averaged *after* applying the sigmoid nonlinearity (similar to what was done in Quang and Xie (2019), except Quang and Xie (2019) used trained conjoined models). This was denoted as “CJ-posthoc-postsigmoid”.

Although we found that the mean performance of RCPS-Avg was higher than the mean performance of RCPS, the 95% confidence intervals were still overlapping. We made a similar observation for CJ-posthoc vs CJ-posthoc-postsigmoid. We did, however, find that CJ-trained significantly outperformed CJ-trained-max on the MAX and SPI1 dataset, consistent with Bartoszewicz *et al.* (2020)’s finding that averaging of representations across the forward and RC strands tended to outperform taking the maximum or the hadamard product. Our findings are displayed in (**Fig. D.9**).

**Figure D.9.**
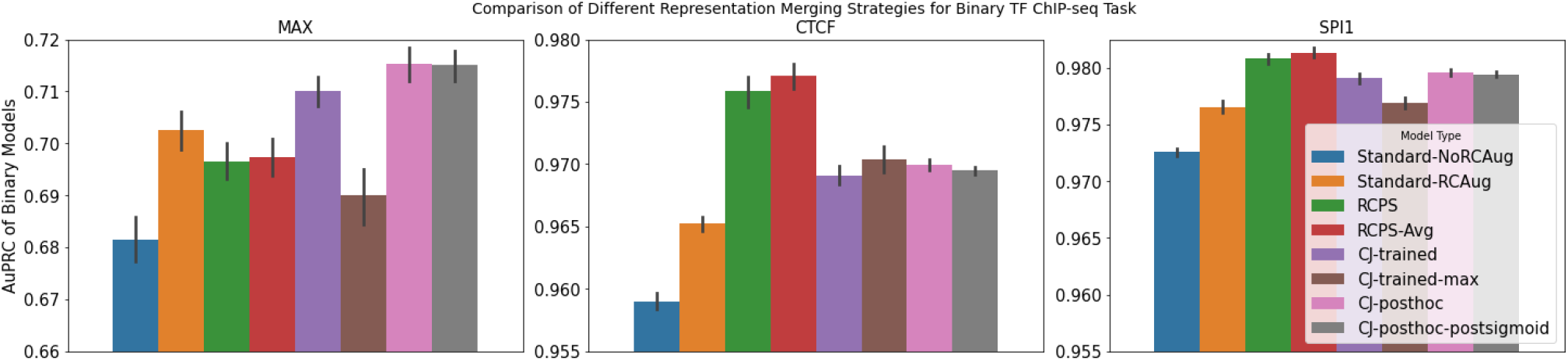
Comparison of different averaging strategies for the conjoined models trained on binary data. Bar heights represent the average performance over 10 random seeds, and error bars represent the 95% confidence intervals for the mean generated by seaborn (Waskom *et al.*, 2017) using 1000 bootstrapped samples. “Standard-RCAug” and “Standard-noRCAug” are standard models trained with and without RC data augmentation. RCPS denotes RC Parameter Sharing. CJ-trained are models that were conjoined during both training and test time. CJ-posthoc are standard models trained with RC data augmentation that were converted to conjoined models only after training. RCPS-Avg is the same as RCPS, but where the strand representations are combined via averaging rather than summing (though averaging is representationally equivalent to summing, it could affect the learning rate). CJ-trained-max is equivalent to CJ-trained, but where the maximum of the sigmoid logits was used rather than the average. CJ-posthoc-sigmoid is equivalent to CJ-posthoc, but but with averaging performed after the sigmoid nonlinearity rather than at the level of the sigmoid logit. Similar to **Fig. 4**, training hyperparameters were set to the tested combination that tended to produced the highest overall AuPRCs.

